# Reconstitution of mammalian Cleavage Factor II involved in 3’ processing of mRNA precursors

**DOI:** 10.1101/366336

**Authors:** Peter Schäfer, Christian Tüting, Lars Schönemann, Uwe Kühn, Thomas Treiber, Nora Treiber, Christian Ihling, Anne Graber, Walter Keller, Gunter Meister, Andrea Sinz, Elmar Wahle

## Abstract

Cleavage factor II (CF II) is a poorly characterized component of the multi-protein complex catalyzing 3’ cleavage and polyadenylation of mammalian mRNA precursors. We have reconstituted CF II as a heterodimer of hPcf11 and hClp1. The heterodimer is active in partially reconstituted cleavage reactions, whereas hClp1 by itself is not. Pcf11 moderately stimulates the RNA 5’ kinase activity of hClp1; the kinase activity is dispensable for RNA cleavage. CF II binds RNA with nanomolar affinity. Binding is mediated mostly by the two zinc fingers in the C-terminal region of hPcf11. RNA is bound without pronounced sequence-specificity, but extended G-rich sequences appear to be preferred. We discuss the possibility that CF II contributes to the recognition of cleavage/polyadenylation substrates through interaction with G-rich far-downstream sequence elements.

## Introduction

Polyadenylated 3’ ends of mRNAs are generated in a two-step processing reaction in which the primary transcript is first cleaved by an endonuclease and the upstream cleavage fragment is then polyadenylated, whereas the downstream fragment is degraded. The reactions are catalyzed, in a tightly coupled manner, by a multi-protein assembly. In mammals, four heterooligomeric complexes are part of this assembly, and many of the subunits contribute to the recognition of the substrate RNA (Xiang et al. 2014; Shi and Manley 2015): Cleavage and polyadenylation specificity factor (CPSF) consists of seven subunits (Shi et al. 2009), among which CPSF30 (= CPSF4) and WDR33 recognize the polyadenylation signal AAUAAA (Chan et al. 2014; Schönemann et al. 2014; Clerici et al. 2017; Sun et al. 2018), and Fip1 binds U-rich sequences (Kaufmann et al. 2004; Lackford et al. 2014). CPSF73 (= CPSF3) cleaves the RNA substrate (Mandel et al. 2006; Eaton et al. 2018). Cleavage stimulation factor (CstF) consists of two copies each of three subunits and binds GU-rich sequence elements downstream of the cleavage site (MacDonald et al. 1994; Beyer et al. 1997; Takagaki and Manley 1997; Martin et al. 2012; Hwang et al. 2016). Cleavage factor I (CF I) contains two copies each of two subunits and associates with UGUA sequences facilitating 3’ processing from a position upstream of AAUAAA (Venkataraman et al. 2005; Yang et al. 2011; Martin et al. 2012; Zhu et al. 2018). Composition and function of cleavage factor II (CF II) will be discussed below. In addition to these complexes, poly(A) polymerase catalyzes poly(A) tail synthesis, and Rbbp6 plays an unknown role in the processing reaction (Di Giammartino et al. 2014). Poly(A) polymerase and a subcomplex of CPSF, mPSF, are sufficient for AAUAAA-dependent polyadenylation of an RNA resembling the cleaved reaction intermediate (Schönemann et al. 2014). In addition, the nuclear poly(A) binding protein (PABPN1) increases the processivity of polyadenylation and contributes to the synthesis of poly(A) tails of a defined length (Kühn et al. 2009; Kühn et al. 2017). This list of 3’ processing factors may well be incomplete.

In addition to the sequence elements listed above, several other blocks of conserved sequences surrounding poly(A) sites have been identified by transcriptome-wide sequence comparisons; these include G-rich motifs located downstream of the CstF binding site (Hu et al. 2005). G-rich downstream elements have also been identified by mutational analyses of the SV40 late poly(A) site (Sadofsky et al. 1985; Qian and Wilusz 1991; Bagga et al. 1995) and other individual poly(A) sites (Yonaha and Proudfoot 1999; Arhin et al. 2002; Oberg et al. 2005; Dalziel et al. 2007).

CF II is the least characterized among the protein complexes constituting the 3’ processing complex. A partial purification suggested that CF II contains hClp1 and hPcf11, orthologues of known 3’ processing factors from *S. cerevisiae* (de Vries et al. 2000). It has remained uncertain whether CF II consists of just these two polypeptides. The MS analysis of a purified mammalian processing complex detected hPcf11 but not hClp1 (Shi et al. 2009). In yeast, a heterodimer of Pcf11 and Clp1 associates with a heterotetramer of Rna14 and Rna15 to form cleavage factor IA (CF IA) (Amrani et al. 1997; Minvielle-Sebastia et al. 1997; Gross and Moore 2001; Gordon et al. 2011; Stojko et al. 2017). The mammalian orthologues of Rna14 and Rna15, CstF77 (= CSTF3) and CstF64 (= CSTF2), are not stably associated with hClp1 and hPcf11 but instead constitute CstF together with a third subunit, CstF50 (= CSTF1), which has no counterpart in yeast (Xiang et al. 2014).

Yeast *clp1* mutants have defects in both steps of the 3’ processing reaction (Haddad et al. 2012). While human and *C. elegans* Clp1 have an RNA 5’ kinase activity (Weitzer and Martinez 2007; Dikfidan et al. 2014), yClp1 is catalytically inactive, suggesting that the activity is not essential for pre-mRNA 3’ processing (Ramirez et al. 2008). This is supported by genetic analysis in mice: Homozygous Clp1 knock-out mice die at an early embryonic stage, but mice homozygous for a kinase-dead Clp1 variant survive for different periods after birth, depending on the genetic background, and have no apparent defect in pre-mRNA 3’ processing (Hanada et al. 2013). Mammalian Clp1 is also part of the tRNA splicing endonuclease, but Pcf11 apparently is not (Paushkin et al. 2004; Weitzer and Martinez 2007). Genetic analyses in mice (Hanada et al. 2013) and humans (Karaca et al. 2014; Schaffer et al. 2014) support a role of Clp1 in tRNA splicing, but how the kinase activity is involved is unclear at the moment (Schaffer et al. 2014; Weitzer et al. 2015).

Mutations in yeast Pcf11 also affect both pre-mRNA cleavage and polyadenylation (Amrani et al. 1997). Yeast Pcf11 is the central subunit of CF IA, interacting with both yClp1 and with the Rna14-Rna15 complex (Gross and Moore 2001; Noble et al. 2007; Gordon et al. 2011). The crystal structure of a complex between yClp1 and a yPcf11 peptide shows interaction surfaces that appear conserved in higher eukaryotes (Noble et al. 2007). Pcf11 is one of the factors responsible for the connection between 3’ end processing and transcription; the protein is specifically required for transcription termination (Birse et al. 1998; Sadowski et al. 2003; Luo et al. 2006; Zhang and Gilmour 2006; West and Proudfoot 2008; Porrua and Libri 2015), and its phosphorylation has been proposed to facilitate transcript release and export (Volanakis et al. 2017). Interestingly, Pcf11, together with Clp1, also participates in the termination of non-coding transcripts that use polyadenylation-independent pathways for 3’ end formation (Kim et al. 2006; Hallais et al. 2013; O’Reilly et al. 2014; Grzechnik et al. 2015; Porrua and Libri 2015). The role of Pcf11 in coupling 3’ end processing to transcription depends on its binding to the serine 2-phosphorylated C-terminal domain (CTD) of RNA polymerase II via a conserved N-terminal CTD interaction domain (CID) (Barilla et al. 2001; Licatalosi et al. 2002; Sadowski et al. 2003; Meinhart and Cramer 2004; Noble et al. 2005; Lunde et al. 2010). A second surface on the ‘body’ of RNA polymerase II may also be involved in the Pcf11 interaction (Pearson and Moore 2014).

Here we present a biochemical reconstitution of human CF II from recombinant hClp1 and hPcf11 and report that CF II contributes to the recognition of 3’ processing substrates, possibly through binding to a G-rich far-downstream element.

## Results

### Human Clp1 and Pcf11 constitute CF II

In order to obtain more definitive information on the subunit composition of CF II, we used a cell line expressing His-FLAG-tagged hClp1 (Paushkin et al. 2004) and isolated the protein by two consecutive affinity purification steps. MS analysis of the main components identified, in addition to hClp1, the tRNA splicing endonuclease complex, as anticipated (Paushkin et al. 2004), and hPcf11 (Fig. 1A). Since the tRNA splicing endonuclease has so far not been directly implicated in 3’ processing, this result suggests that CF II may consist of just hPcf11 and hClp1. These two polypeptides, with an N-terminal his-tag on hClp1, were therefore co-expressed in insect cells by means of the MultiBac system (Berger et al. 2004; Fitzgerald et al. 2006). Chromatography on Ni^2+^ beads resulted in the co-purification of both proteins, which remained associated during subsequent anion exchange chromatography and gel filtration (Fig. 1B, C, E, F). Identity and completeness of both polypeptides were confirmed by LC/MS/MS. In gel filtration, CF II eluted ahead of the largest standard protein with an extrapolated Stoke’s radius of 7.1 nm (Fig. 1E). Glycerol gradient centrifugation indicated a sedimentation coefficient of 7S (data not shown). A native molecular weight of 207 kDa calculated from these data (Siegel and Monty 1966; Erickson 2009) is in good agreement with the theoretical molecular weight of a heterodimeric hPcf11•hClp1 complex (220.8 kDa), consistent with the stoichiometry of the orthologues in yeast CF IA (Gordon et al. 2011; Stojko et al. 2017).

**Fig. 1:**
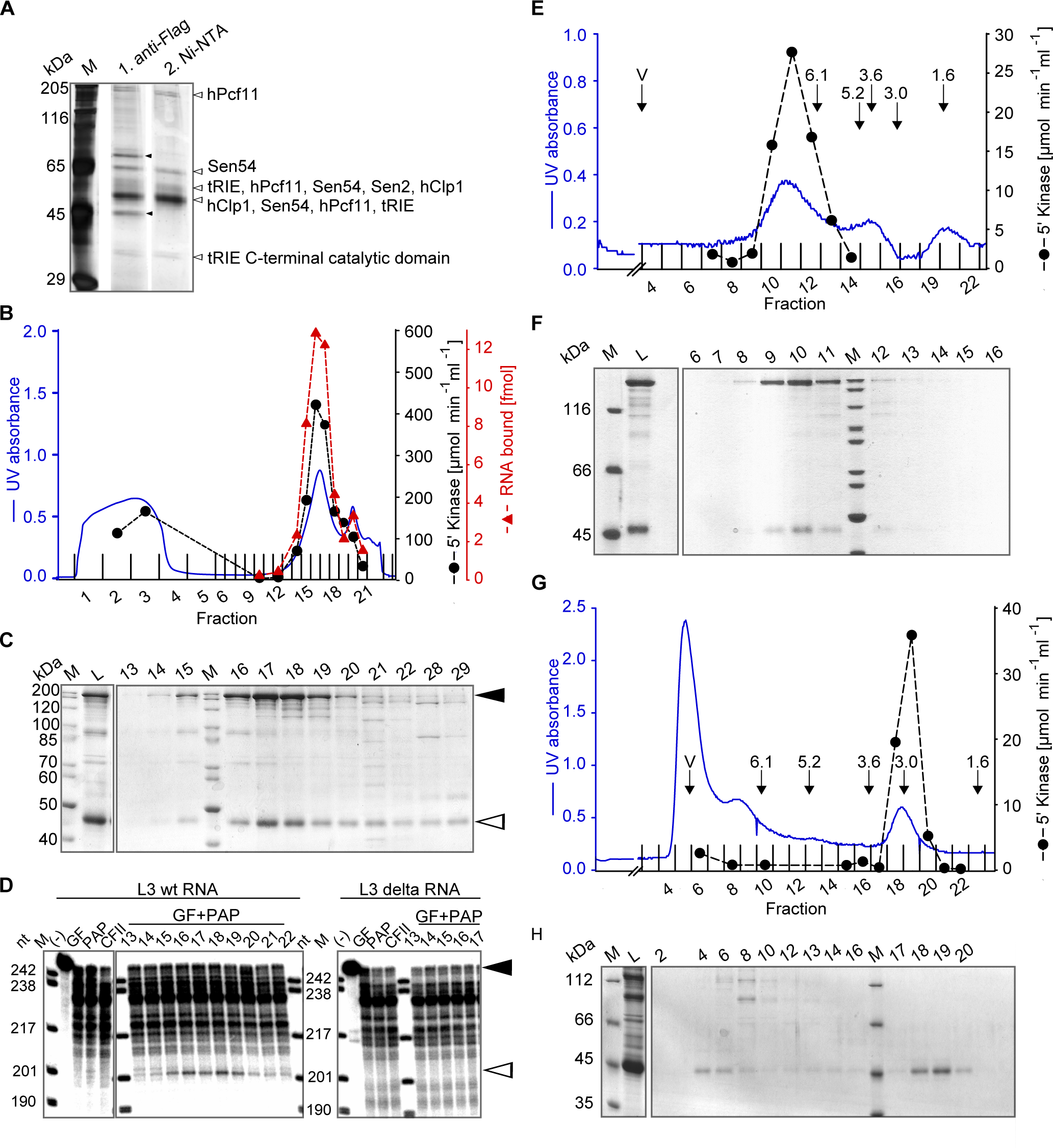
CF II is a heterodimer of hClp1 and hPcf11 and co-purifies with RNA 5’ kinase, RNA binding and 3’ cleavage activity. (A) His-FLAG-tagged hClp1 was affinity purified from a stable cell line by two consecutive purification steps on FLAG beads and Ni^2+^ beads. The silver-stained SDS gel shows the eluates from the first (middle lane) and second purification step (right lane). The two bands marked in the FLAG eluate are PRMT5 and MEP50, known FLAG-binding proteins. Sen2, Sen54 and tRIE are subunits of the tRNA splicing endonuclease. (B) MonoQ column profile of his-tagged hClp1 and hPcf11 co-expressed in baculovirus-infected insect cells (see Material and Methods). RNA binding was measured by nitrocellulose filter binding assays with 0.5 µl of a 1:100 dilution of the fractions indicated and 100 fmol of SV40 late RNA (see Materials and Methods). Kinase activity was assayed as described in Materials and Methods. (C) Fractions of the column shown in (B) were analyzed by SDS-polyacrylamide gel electrophoresis. Arrowheads indicate hPcf11 and hClp1. L, load of column. (D) Fractions of the column shown in (B) were assayed for reconstitution of 3’ cleavage as described in Materials and Methods. Assays contained poly(A) polymerase (PAP) and a fraction (GF) generated by ammonium sulfate precipitation of HeLa cell nuclear extract and Superose 6 gel filtration; this fraction contained all factors required for cleavage except PAP and CF II. Reactions were complemented with 2 µl of the MonoQ column fractions and were carried out with wild-type L3 RNA or L3Δ RNA with a point mutation in the AAUAAA signal. With both RNAs, the first four lanes following the marker lane show negative controls, as indicated. Black and white arrowheads indicate RNA substrates and specific cleavage products. AAUAAA- and CF II-independent partial degradation of most of the substrate is due to nuclease activity in GF (compare control lanes). (E) Gel filtration of CF II on a Superose 6 column. Arrows indicate the void volume (V) and the elution peaks of marker proteins with their Stoke’s radii (see Materials and Methods). (F) SDS polyacrylamide gel electrophoresis of the column fractions of (E). (G) Superdex 200 gel filtration column profile of His-tagged hClp1 expressed in baculovirus-infected cells (see Materials and Methods). RNA kinase activity was measured as described in Materials and Methods. The void volume (V) and peak positions of marker proteins with their Stoke’s radii are indicated. (H) SDS polyacrylamide gel electrophoresis of the column fractions of (G).

His-tagged hClp1 was also expressed by itself and purified by metal affinity chromatography. Subsequent gel filtration showed an apparent native molecular weight near 40 kDa, consistent with the calculated molecular weight of 48.7 kDa and, thus, a monomeric structure (Fig. 1G, H). Human Pcf11 expressed in the absence of hClp1 was insoluble.

The hClp1•hPcf11 complex supported AAUAAA-dependent 3’ cleavage in partially reconstituted assays (Fig. 2C). In these assays, replacement of ATP by cordycepin triphosphate (3’-dATP) prevented polyadenylation and permitted a direct detection of the upstream cleavage fragment. The cleavage activity co-purified with the hClp1•hPcf11 complex on the MonoQ column (Fig. 1D). In contrast, hClp1 did not support cleavage (Fig. 2C). We conclude that hClp1 and hPcf11 constitute functional CF II.

**Fig. 2:**
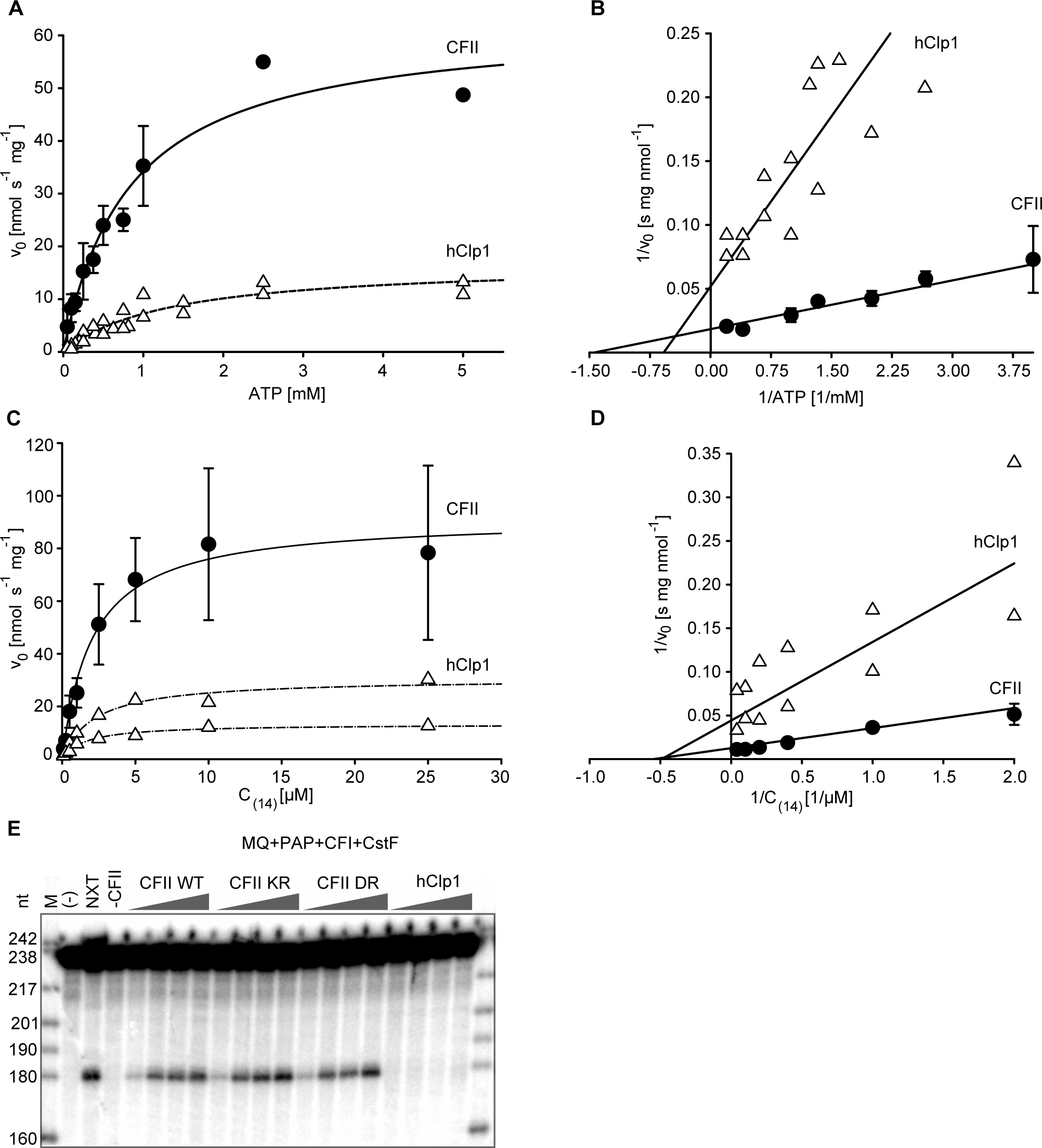
CF II has RNA 5’ kinase activity, which is not required for RNA cleavage. (A) Initial velocities of the 5’ kinase activities of CF II or hClp1 were measured at varying concentrations of [γ-^32^P]-ATP and a constant concentration of C_14_ as described in Materials and Methods. Plots represent fits to the Michaelis-Menten equation. Data points with error bars representing the standard deviation were averaged from n ≥3. Titrations for hClp1 were done twice, and data from both experiments were combined into one fit. (B) Data from (A) are presented as a Lineweaver-Burk (double-reciprocal) plot. Fits are based on all data points, but only data for higher ATP concentrations are shown so that the intersections with the axes can be seen more easily. (C) Initial velocities of the 5’ kinase activities of CF II or hClp1 were measured at a constant concentration of [γ-^32^P]-ATP and varying concentrations of C_14_. Plots were as in (A). Titrations for hClp1 were done twice and fitted separately. (D) Data from (C) are presented as a Lineweaver-Burk (double-reciprocal) plot. The fit for hClp1 is based on both titrations. Fits are based on all data points, but only data for higher RNA concentrations are shown so that the intersections with the axes can be seen more easily. In both ATP and RNA titrations, the apparently higher V_max_ values of CF II as opposed to hClp1 have to be viewed with caution as discussed in the legend to Table 1. (E) Cleavage assays contained 5, 25, 50 or 200 fmol of hClp1 or of wild-type CF II or of CF II with clustered point mutations in the active site of hCp1 (KR: K127A, R288A, R293L; DR: D151A, R288A, R293L). These proteins were tested in a complementation system containing the SV40 late RNA, poly(A) polymerase, CF I, CstF and a protein fraction generated from HeLa cell nuclear extract by ammonium sulfate precipitation, gel filtration and MonoQ chromatography as described in Materials and Methods. Note that the MonoQ fraction contained less unspecific nuclease activity than the gel filtration fraction used in Fig. 1D. Omission of CF II or all proteins was used as negative controls and addition of nuclear extract (NXT) as a positive control, as indicated. Black and white arrowheads indicate substrate RNA and 5’ cleavage product, respectively. Controls with a mutant RNA showed that the cleavage activities were dependent on the AAUAAA sequence (data not shown).

### CF II has RNA 5’ kinase activity, which is not essential for 3’ processing

The anticipated RNA 5’ kinase activity of hClp1 was also manifest in CF II (Fig. 1B, E). Steady-state kinetics revealed that CF II had a moderately reduced K_M_ for ATP as compared to hClp1 (Fig. 2A, B; Table 1). The K_M_ for oligocytidine (C_14_) was not noticeably affected (Fig. 2C, D; Table 1). Measured k_cat_ values for CF II were higher than those for hClp1, but this may have been due to technical reasons (see legend to Table 1). Also, hClp1 activity declined during storage whereas CF II was stable.

**Table 1:** Steady-state parameters for the RNA 5’ kinase activity of CFII and hClp1

| Substrate | Protein | $k_{cat}$ [$s^{-1}$] | $K_M$ [$\mu M$] | $k_{cat}/K_M$ [$1/s \cdot M$] |
| --- | --- | --- | --- | --- |
| ATP | CF II | $2.8 \pm 0.8$ | $700 \pm 200$ | $4300 \pm 1400$ |
| | hClp1 | $1.3 \pm 0.5$ | $2500 \pm 1100$ | $500 \pm 100$ |
| $C_{14}$ | CF II | $4.5 \pm 1.4$ | $2.5 \pm 0.4$ | $1.8 \times 10^6 \pm 0.4 \times 10^6$ |
| | hClp1 | ($1.6/0.7$) | $2.4/1.6$ | ($0.7 \times 10^6/0.4 \times 10^6$) |

Kinase assays were carried out as described in Materials and Methods. $K_M$ and $V_{max}$ were obtained from standard Michaelis-Menten analysis, and $V_{max}$ values were converted to $k_{cat}$ . The $C_{14}$ titration with hClp1 was carried out twice, and the parameters derived from the two experiments are both listed. All other numbers listed represent the average ± standard deviation ( $n = 3-4$ ). The $k_{cat}$ values for hClp1 and the resulting $k_{cat}/K_M$ values are put in brackets because the activity of hClp1 declined upon storage; CF II, in contrast, was stable. This contributes to a lower apparent $k_{cat}$ of hClp1 but should not affect $K_M$ determinations. Note also that the ATP concentration used for $C_{14}$ titrations was $1.2 \times K_M$ for CF II but $0.75 \times K_M$ for hClp1; saturating concentrations would have required prohibitively large amounts of radioactivity. Finally, different assays were used to measure kinase activity in ATP versus $C_{14}$ titrations (see Materials and Methods).

A single point mutation, K127A, introduced into the conserved active site of hClp1 (Dikfidan et al. 2014), reduced the kinase activity of CFII by only a factor of 30. However, triple mutations K127A, R288A, R293L (abbreviated KR) or D151A, R288A, R293L (abbreviated DR) reduced the activity to undetectable levels, less than 0.2% of the wild-type (data not shown). All mutant proteins could be co-purified in a complex with hPcf11 (data not shown). CF II containing the kinase-dead Clp1 variants was active in 3’ cleavage (Fig. 2E). Thus, the kinase activity is not essential for 3’ processing.

### Domain organization of CF II

In contrast to Clp1 (Noble et al. 2007; Dikfidan et al. 2014), Pcf11 is structurally not well characterized. A scheme of the protein is shown in Fig. 3A: The N-terminal CID (Meinhart and Cramer 2004) is followed by a helical domain of unknown function (Xu et al. 2015). Amino acids 295 - 565 comprise a highly charged region with 28.4 % basic and 14.8% acidic residues and a serine content of 14.8%. Amino acids 770 - 1123 contain 30 repeats of a motif of about thirteen amino acids (FEGP repeats for short) with a high content of proline and glycine (Fig. 3B). Between 22 and 36 repeats of the FEGP motif are conserved in Pcf11 orthologues in vertebrates, but not in *Drosophila*, *C. elegans* or yeast. Data base searches did not reveal similar repeats in other proteins. Most arginine residues in the FEGP repeats are asymmetrically dimethylated (Guo et al. 2014). Numerous dimethylarginine residues in the repeats were also detected in the MS analysis of our baculovirus-produced hPcf11 (data not shown). The C-terminus of Pcf11 contains two zinc fingers (Barilla et al. 2001; Sadowski et al. 2003; Guegueniat et al. 2017; Yang et al. 2017), which straddle the Clp1 interaction site (Noble et al. 2007). The region of yPcf11 responsible for the Rna14/Rna15 interaction is on the N-terminal side of the first zinc finger (Amrani et al. 1997) and has been narrowed down to aa 331-417 (Lionel Minvielle-Sebastia, personal communication). This region has no obvious counterpart in hPcf11.

**Fig. 3:**
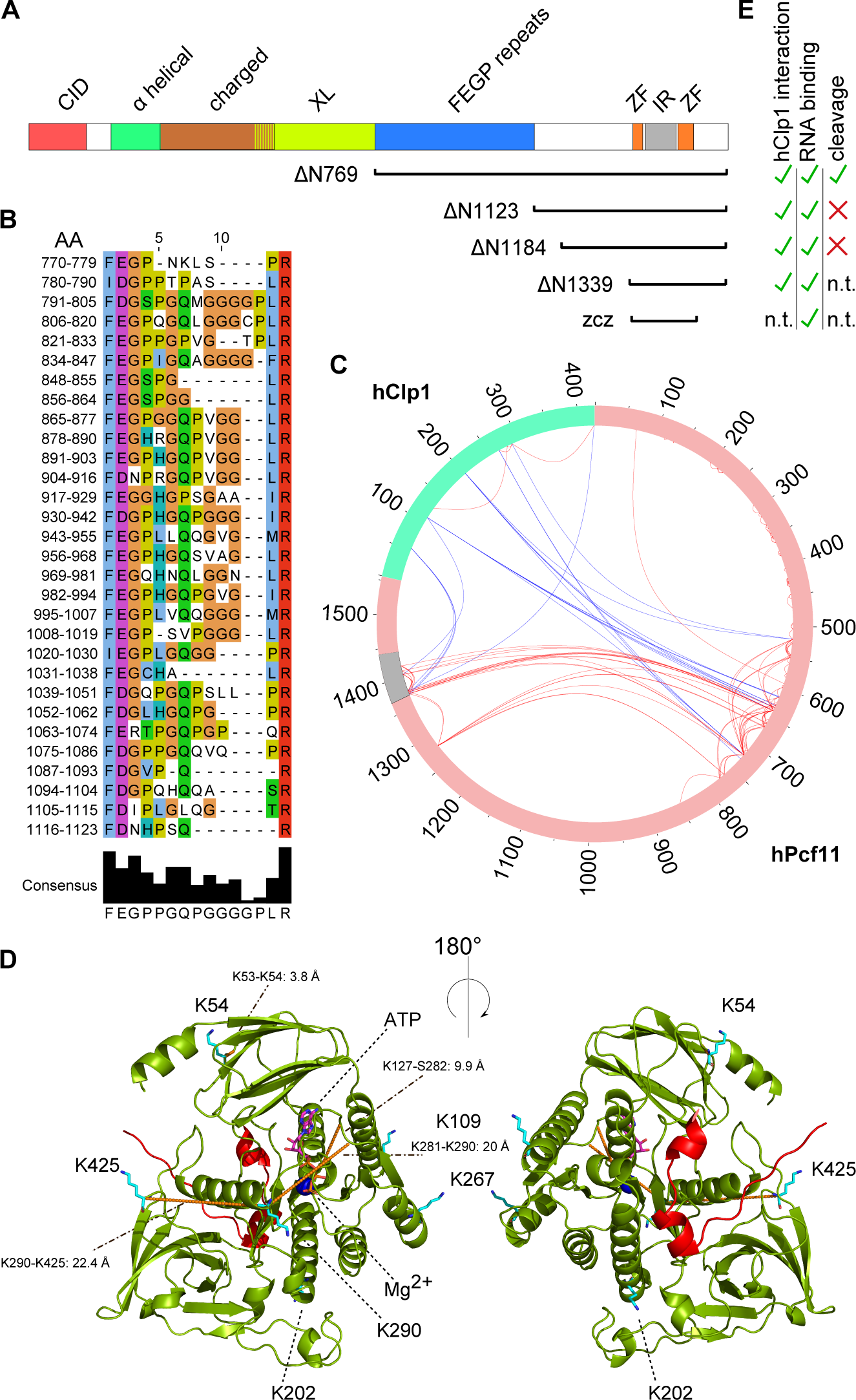
Domain organization and cross-linking of CF II. (A) Domains of hPcf11. Highlighted are the CID (CTD Interaction Domain; residues 14-142), an α-helical region (183-297), a highly charged, basic region (295 - 550) overlapping with the region cross-linked to hClp1 (XL; 500-770), the FEGP repeats (770-1124), the zinc fingers (ZF; 1343-1368 and 1443-1478) and the hClp1 interaction region (IR; 1371-1439). The bars at the bottom indicate parts of hPcf11 present in the deletion variants. (B) Alignment of consecutive residues 770 - 1126 of hPcf11 reveals FEGP repeats. The alignment was produced with JalView (Waterhouse et al. 2009). (C) Overview of cross-links in CF II. The circle represents the primary structures of hClp1 and hPcf11 with amino acid numbering indicated. Blue lines indicate intermolecular and red lines intramolecular cross-links. The Clp1 interaction region of Pcf11 based on the crystal structure of the yeast proteins is in grey. The K53-K54 cross-link in hClp1 is not visible in the plot. (D) Cross-link sites in a model of hClp1 (green) with the superimposed peptide from yPcf11 (red) (PDB accession number: 4OI4). Lysine side-chains forming inter- or intramolecular cross-links are shown. Intramolecular cross-links are shown in orange with participating side-chains and C_α-_C_α-_distances (in Ångstrom) indicated. The structure of hClp1 was modeled with the Robetta Server (http://robetta.bakerlab.org) (Kim et al. 2004).. (E) Summary of the activities attributed to the hPcf11 deletion variants (n. t., not tested).

The interaction between hPcf11 and hClp1 was analyzed by treatment of the complex with the MS-cleavable DSBU cross-linker that covalently links amino groups and, with a lower efficiency, hydroxy groups at a maximum Cα-Cα distance of ~ 30 Å (Müller et al. 2010). Cross-linked proteins were digested with trypsin and peptides analyzed by LC/MS/MS (Suppl. Table 1). The analysis revealed four intramolecular cross-links of hClp1. All were consistent, within the 30 Å distance, with a three-dimensional homology model of the protein based on the structures of the *S. cerevisiae* and *C. elegans* orthologues (Fig. 3C, D). The analysis also revealed eighteen intermolecular cross-links between hClp1 and hPCF11 (Fig. 3C). Five cross-links from Clp1 residues 54, 109 and 425 to the region 1371 - 1405 of hPcf11 (within the box labeled IR in Fig. 3A) were consistent with the interaction surfaces of the yeast orthologues (Noble et al. 2007) (Fig. 3D). Thus, the interaction between Clp1 and Pcf11 is conserved. Lysine residues 109, 202, 267 and 290 of hClp1 formed numerous cross-links to a region of hPcf11 between residues 520 and 731 (labeled XL in Fig. 3A) (Fig. 3C, D). This region also contained multiple intramolecular cross-links between amino acids widely separated in the primary structure, suggesting the possibility of a globular fold. The two separate regions of hPcf11 that were cross-linked to hClp1 were also cross-linked with each other, indicating their neighborhood in the quaternary structure of CF II.

Limited trypsin digestion of CF II under native conditions resulted in rapid degradation of hPcf11, with accumulation of a stable fragment that migrated at a little over 50 KDa in SDS gels (Fig. 4A). Analysis by MS/MS and N-terminal Edman degradation suggested Ala1185 as the N-terminus of the trypsin-resistant fragment. The fragment remained associated with his-tagged hClp1 at the end of trypsin digestion. The corresponding engineered fragment hPcf11 ΔN1184 was also associated with hClp1 upon co-expression (data not shown). Even a short C-terminal hPcf11 fragment starting with the first zinc finger (ΔN1339) co-purified with hClp1 when the two were co-expressed in insect cells (Fig. 4B). Thus, the conserved interaction surface between the zinc fingers is sufficient for a stable interaction; the more N-terminal region of hPcf11 that is cross-linked to hClp1 is not required.

**Fig. 4:**
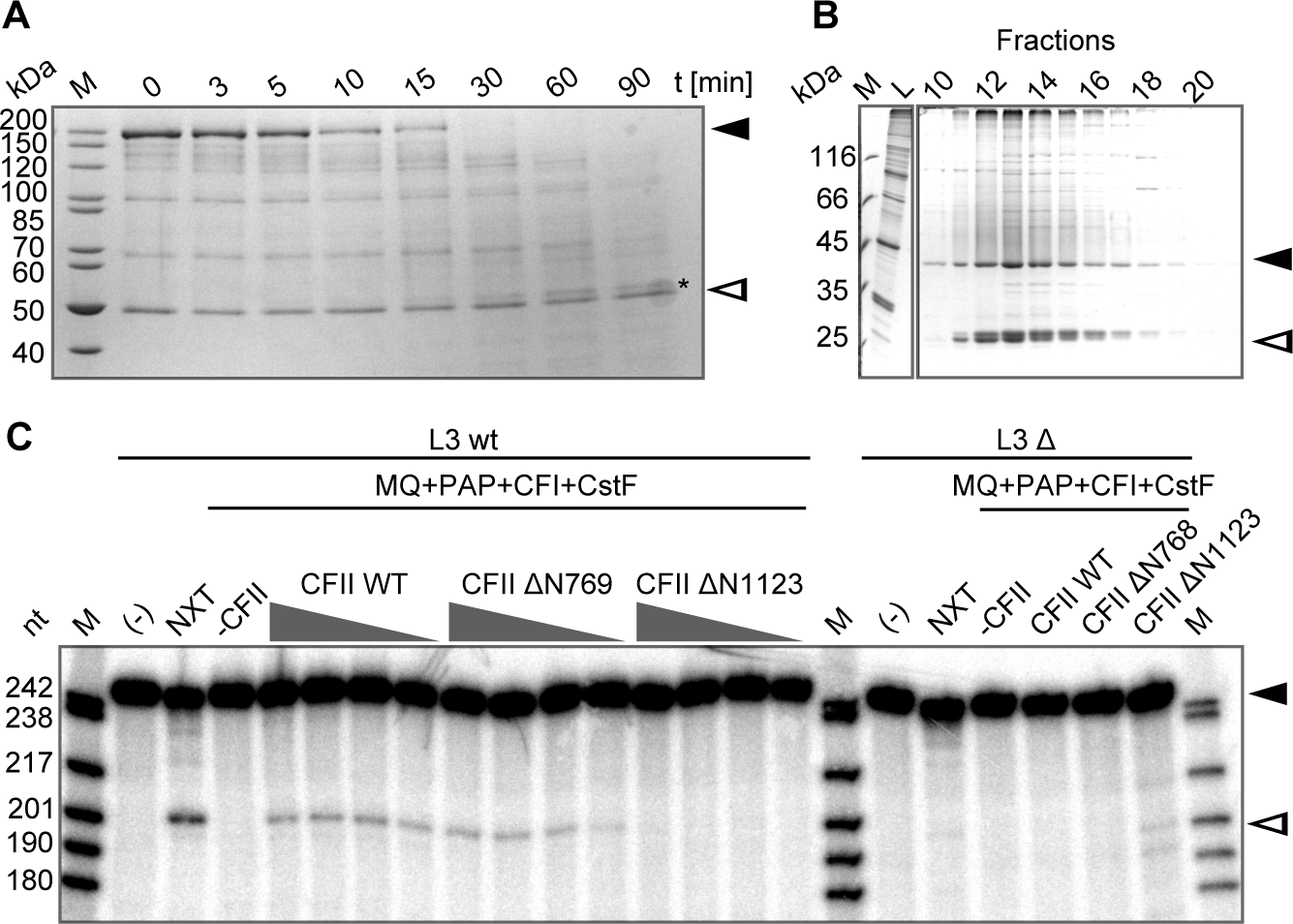
Domains of hPcf11. (A) Trypsin digestion identifies a stable C-terminal hPcf11 fragment. Human Pcf11 was digested with trypsin for the times indicated (see Materials and Methods). Proteins were analyzed on an SDS-polyacrylamide gel and stained with Coomassie. Black and white arrowheads indicate full-length hPcf11 and hClp1, respectively, and the asterisk marks the stable hPcf11 fragment. (B) The C-terminal region of hPcf11 containing the hClp1 interaction sequence flanked by the zinc fingers suffices for stable hClp1 binding. Untagged hClp1 and his-tagged hPcf11ΔN1339 were co-expressed and purified via Ni-NTA and MonoQ chromatography. The figure shows an analysis of the MonoQ peak fractions by SDS-polyacrylamide gel electrophoresis and Coomassie staining. Black and white arrowheads indicate hClp1 and the hPcf11 fragment, respectively. Identity and completeness of the hPcf11 fragment were confirmed by LC/MS/MS. (C) A C-terminal hPcf11 fragment starting with the FEGP repeats supports pre-mRNA cleavage. Cleavage assays were performed with the L3 RNA and mutant control and proteins as described in the legend to Fig. 2B. Controls were as in Fig. 2B. CF II ΔN769 and ΔN1123 refers to the heterodimers reconstituted with the respective hPcf11 mutants. Black and white arrowheads indicate substrate RNA and 5’ cleavage product, respectively.

Several additional deletion variants of hPcf11 were co-expressed with hClp1 and purified for use in cleavage assays (Fig. 3A, E). ΔN769, starting with the FEGP repeats, was the largest fragment tested and the only one providing CF II function. In contrast, the next smaller fragment ΔN1123, starting just C-terminal of the FEGP repeats, was inactive (Fig. 4C). Both hPcf11 variants were co-purified with hClp1, and the complexes had similar activities in RNA binding assays (see below); thus, the inactive variant is not misfolded. A CF II complex containing hPCF11 ΔN1184 was also inactive in cleavage (Fig. 3E; data not shown). We conclude that the FEGP repeats are essential, but everything N-terminal of the repeats is dispensable for RNA cleavage *in vitro*. This includes the CID, in agreement with data obtained in yeast (Sadowski et al. 2003), and the highly charged region.

The stable association of yPcf11 and yClp1 with Rna14 and RNA15 suggests that CF II might interact with CstF (see Introduction). However, when CF II and CstF were mixed at a final concentration of 0.4 µM each and analyzed by gel filtration, no association was detected. Likewise, gel filtration did not detect an interaction between CF II and the C-terminal alpha-helical bundle of CstF-64 (purified as a His-Sumo fusion) (Qu et al. 2007). Gel filtration of a mixture of CF I and CF II at 0.7 µM each did not reveal an association either, and no interaction was found when CstF was added as a third component. Pull-down experiments in which FLAG-tagged CF II containing hPcf11 ΔN769 was incubated with CF I or CstF were likewise negative. CF II is an RNA binding protein (see below), and even weak interactions with other RNA binding proteins, not detectable by gel filtration or pull-down experiments, should be visible as cooperative RNA binding. However, when CF I, CF II and CstF were used in filter-binding experiments at concentrations below the K_D_, mixing of the proteins in all pairwise combinations resulted in additive binding, i. e. no cooperativity was visible.

### CF II binds RNA via the zinc fingers of hPcf11

Nitrocellulose filter-binding experiments revealed that CF II avidly bound to RNA: An RNA of 230 nt containing the SV40 late 3’ processing site was bound with a K_50_ of ~ 0.5 nM. The adenovirus L3 3’ processing site was bound with comparable affinity (Fig. 5A). The RNA binding activity co-migrated with CF II in anion exchange chromatography (Fig. 1B) and gel filtration (data not shown).

**Fig. 5:**
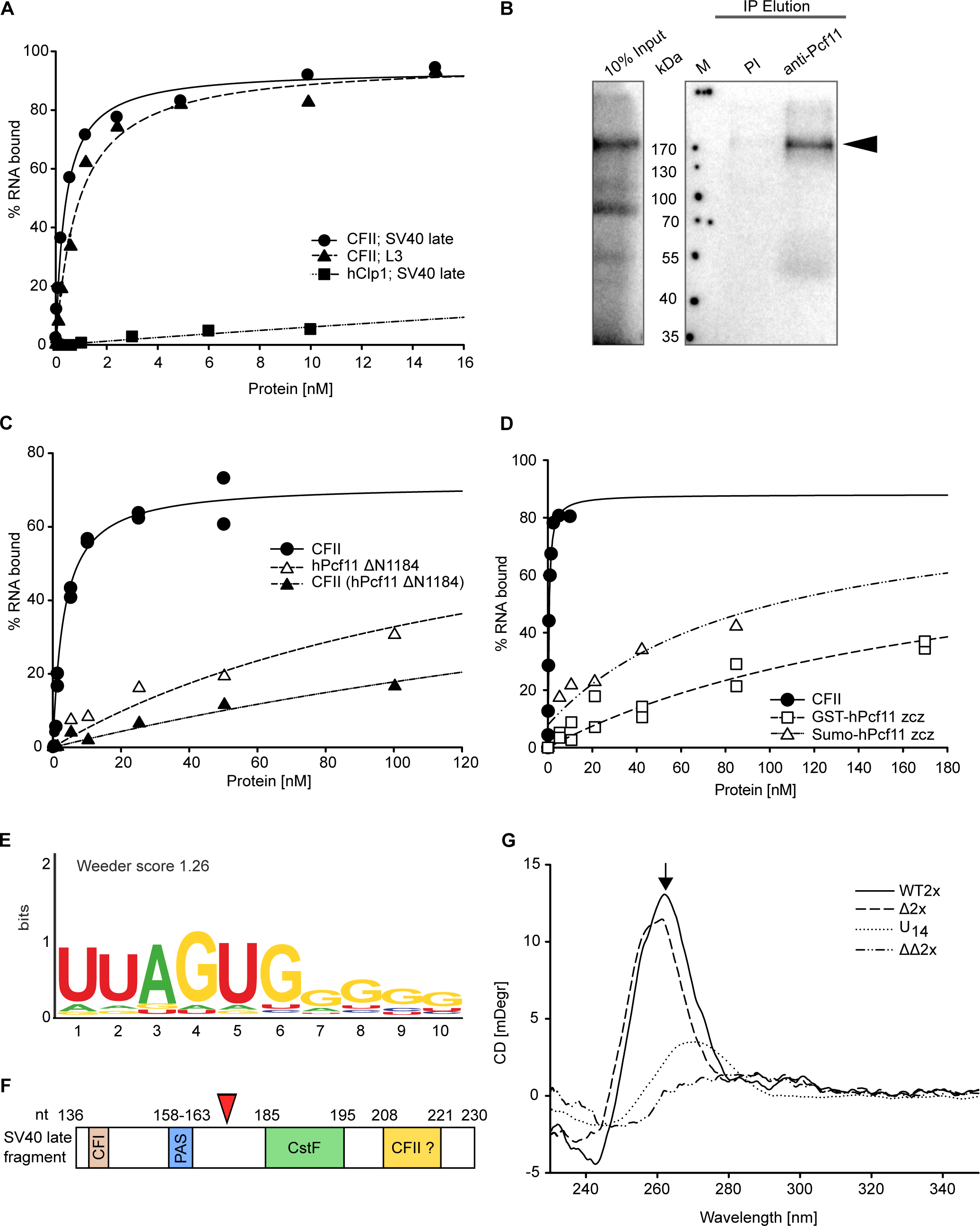
CF II binds RNA via the zinc fingers of Pcf11. (A) Nitrocellulose filter-binding experiments showing binding of CF II or hClp1 to SV40 late or L3 RNAs, as indicated. Hyperbolic fitting indicated K_50_ values of 0.4 nM and 0.9 nM for CF II binding to the SV40 late or the L3 RNA, respectively. The extrapolated K_50_ for hClp1 was ~ 140 nM. However, even this relatively low RNA binding activity was at least partially due to a contamination, as indicated by analysis of the profile of the gel filtration column from which hClp1 was obtained. (B) CF II was incubated with 5’-labeled U_14_ and irradiated with UV light. An aliquot of the cross-linked material was then subjected to immunoprecipitation with antibodies against hPcf11 or pre-immune serum as a control. Input and precipitated material were analyzed by SDS-polyacrylamide gel electrophoresis and phosphoimaging. (C) Gel shift experiments with the RNA oligonucleotide 208-221 (Table 2) comparing wild-type CF II, CF II reconstituted with hPcf11ΔN1184, and hPcf11ΔN1184 by itself. Fits indicated K_50_ values of 3.3 nM for CF II and >100 nM for the two other proteins. (D) Binding of HisSumo-hPcf11zcz (single titration) and GST-hPcf11zcz (n = 2) to RNA oligonucleotide 191-230 (Table 2). Extrapolated K_50_ values were ~ 100 and 200 nM. For comparison, binding of the same RNA to CF II is shown (K_50_ = 0.5 nM). (E) Sequence logo derived from a selection carried out with hPcf11ΔN1184 and nitrocellulose filter binding. (F) A scheme of the SV40 late 3’ processing signal. CFI and and CstF indicate the respective protein binding sites, PAS is the AAUAAA signal. The yellow box represents the G-rich sequence described in the Introduction. The cleavage site (red triangle) is after nucleotide 176. Numbering of the sequence is as in Table 2. (G) CD spectra of RNA oligonucleotides wt2x and Δ2x show a maximum at 260 nm, indicating a G quadruplex structure, whereas ΔΔ2x has a spectrum similar to that of unstructured U_14_. The 260 nm peak of Δ2x was sensitive to heating to 90°C.

RNA binding by hClp1 was at least 300fold weaker compared to CF II and could not unequivocally be attributed to hClp1 as opposed to a potential contaminant of the protein preparation (Fig. 5A and data not shown). Thus, hPcf11 is important for the RNA binding activity of CF II. UV cross-linking followed by immunoprecipitation confirmed RNA binding to hPcf11 (Fig. 5B). So far, two RNA binding domains have been described in yPcf11: The CID weakly binds RNA (Zhang et al. 2005; Hollingworth et al. 2006), and the zinc finger domain has also been shown, by UV cross-linking, to interact with RNA. No sequence specificity was detected (Guegueniat et al. 2017). In gel shift assays with several RNAs, CF II containing full-length hPcf11 bound much more tightly than CF II containing N-terminally truncated hPcf11 variants (Fig. 5C and data not shown). These data are consistent with a contribution of the CID to RNA binding. The hPcf11 ΔN1184 fragment by itself bound with a similar affinity as the heterodimer with hClp1 (Fig. 5C), confirming that hClp1 does not significantly contribute to RNA binding.

**Table 2:**
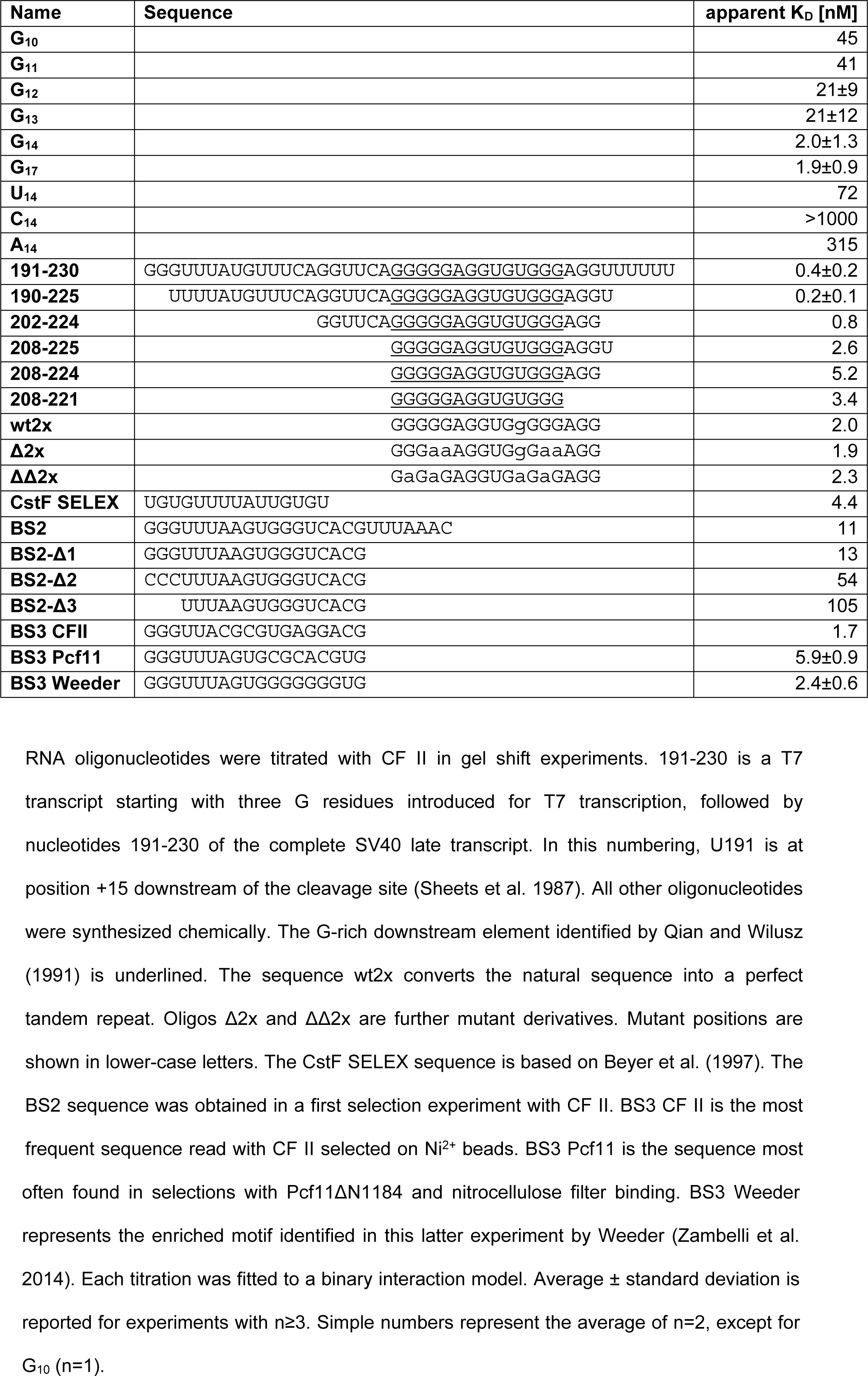
Affinities of CF II for RNA oligonucleotides

In order to assess the role of the zinc fingers in RNA binding, we expressed and purified an hPcf11 fragment containing just the zinc fingers and the intervening Clp1 interaction sequence (zcz fragment; amino acids 1340 - 1505), fused to either His-Sumo or GST. Both fusion proteins, purified by different procedures, bound RNA in filter-binding and gel shift assays. The RNA binding activity co-migrated with the fusion proteins in anion exchange chromatography (data not shown). The RNA 191-230 was bound with an apparent K_D_ of 100 - 200 nM (Fig. 5D), approximately 200fold higher than the K_D_ of CF II (Table 2; see below). Thus, approximately 75% of the free energy of binding is provided by the two zinc fingers. The GST-zcz fusion protein bound G_14_ in gel shift assays but not the other homopolymeric 14mers. Thus, the RNA binding specificity of the zinc fingers matches the specificity of CF II (see below).

The results of the hPcf11 deletion analyses are summarized in Fig. 3E.

### CF II prefers G-rich sequences

In gel shift assays, CF II bound G_14_ with high affinity (~ 2 nM), whereas the other three homopolymeric 14mers were bound weakly or very weakly. G_17_ was bound as tightly as G_14_, but shorter G homopolymers were bound less well (Table 2). Thus, CF II appears to have an extended binding site of ~14 nucleotides.

PAR-CLIP experiments have mapped a peak of yPcf11 binding to pre-mRNAs 130 nt downstream of the cleavage site (Baejen et al. 2017); no specific sequence motif was identified. ChIP experiments also showed binding of *S. cerevisiae* and *S. pombe* Pcf11 downstream of the cleavage site (Mayer et al. 2012; Baejen et al. 2017; Wittmann et al. 2017). Based on this position of binding and the preference of CF II for oligo(G), we tested whether CF II might be responsible for the recognition of the G-rich downstream element mapped in the SV40 late polyadenylation signal (see Introduction; Fig. 5F). Indeed, a series of oligonucleotides gradually shortened to the G-rich motif (208-221) revealed tight binding (Table 2). However, two variant sequences with reduced G content (from 13/17 in 208-224 to 10/17 in Δ2x and ΔΔ2x; Table 2) had similar affinities. Even a 16mer isolated as a good ligand for CstF (CstF selex; Table 2) was bound well, although it was not G-rich.

The binding specificity of CF II was further investigated by selection of binders from a random pool (’bind-n-seq) (Lambert et al. 2015). A pool containing synthetic RNA 14mers followed by an invariant GUUU sequence was used in a first selection round. The selected RNA was cloned, and the presence of a T7 promoter in the cloning adaptor allowed for subsequent transcription for a second round of selection. T7 transcription resulted in the addition of a GGG sequence at the 5’ end. Sequences resulting from a first selection experiment were synthesized and tested in gel shift assays (BS2 sequences; Table 2). Only moderate affinities were observed, but variation of the selected sequence showed that the 5’GGG sequence improved CF II binding by five- to tenfold, supporting the importance of G residues for CF II binding. In contrast, the three invariant U residues at the 3’ end were irrelevant. Therefore, winner sequences from subsequent selection experiments were synthesized with the 5’ GGG extension but without the 3’ UUU sequence (BS3 sequences; Table 2). In one type of experiment, RNA was selected with CF II and binding to Ni^2+^ beads. In both replicates of this experiment, multiple reads of the sequence BS3 CF II (Table 2), with some variation in position 6, were found; these sequences were not present in the no-protein control. In the second type of experiment, selection with hPcf11 ΔN1184 and nitrocellulose filter binding, sequences related to BS3 Pcf11 were enriched in both replicates. In this case, statistical analysis (Zambelli et al. 2014) identified the sequence BS3 Weeder (Fig. 5E). All three winner sequences tested were G-rich and bound CF II with high affinity (Table 2). Thus, the results support a preference of CF II for G residues.

Many of the G-rich oligonucleotides bound by CF II are predicted to form intramolecular G-quadruplex (G4) structures (Bedrat et al. 2016). Indeed, CD spectroscopy of selected oligonucleotides confirmed G4 formation (Fig. 5G). However, two results suggest that CF II binds its G-rich ligands in a non-G4 form: First, whereas G4 formation leads to an accelerated migration of RNA during native gel electrophoresis (Wang et al. 2017), the oligonucleotides listed in Table 2 migrated as expected from their lengths. Thus, we assume that the G4 structures detected by CD spectroscopy were intermolecular structures favored by high RNA concentrations (2 µM), whereas no structures were formed at the low concentrations (0.1 nM) used in gel shift assays. Second, both the oligonucleotide wt2x, derived from the SV40 late sequence 208-224 by a single mutation that generates a direct repeat of 8 nucleotides, and a mutant derivative, Δ2x, contain four blocks of G_≥2_ and are thus predicted to form G4 structures. In contrast, oligonucleotide ΔΔ2x has the same nucleotide composition as Δ2x but only two G_2_ blocks; thus, it should not form G4 structures. CD spectroscopy confirmed (presumably intermolecular) G4 formation by the wt2x and Δ2x oligonucleotides but not by ΔΔ2x (Fig. 5G). Still, CF II bound all three oligonucleotides with similarly high affinities (Table 2). Thus, all three RNAs are presumably bound in non-G4 conformations.

The association of the high-affinity RNA binder hPcf11 with the RNA kinase hClp1 would suggest that ligands preferred by hPcf11 should also be better substrates for hClp1. However, in a comparison of two oligonucleotides, C_14_ (poor binder) and 190-225 (good binder) (Table 2), no such difference was observed. The reason for this result is currently unknown.

## Discussion

Among the protein complexes contributing to 3’ processing of mammalian mRNA precursors, CstF and CF I have previously been reconstituted from recombinant proteins (Takagaki and Manley 1994; Rüegsegger et al. 1998; Yang et al. 2011; Yang et al. 2018). Of the central factor, CPSF, only the mPSF subcomplex, active in polyadenylation, has been reconstituted (Schönemann et al. 2014; Clerici et al. 2017; Clerici et al. 2018; Sun et al. 2018), but the subunit composition of CPSF is reasonably clear from biochemical experiments (Bienroth et al. 1991; Shi et al. 2009; Schönemann et al. 2014). In contrast, CF II had never been purified to the point that its subunit composition could be judged. Here we show by affinity purification of a tagged subunit and by overexpression in a heterologous system that CF II consists of just two polypeptides, hPcf11 and hClp1. These two proteins form a heterodimer, which is active in a partially reconstituted 3’ processing reaction.

Human Clp1 and hPcf11 interact via conserved surfaces; this was predicted by X-ray crystallography of the yeast complex and sequence conservation in the human proteins (Noble et al. 2007) and was confirmed here by cross-linking and MS analysis. The hPcf11 subunit is essential for 3’ processing; hClp1 by itself is inactive. Within hPcf11, the N-terminal CID is dispensable, in agreement with data in yeast (Sadowski et al. 2003). In contrast, a unique repeat structure, the FEGP repeats, is necessary for CF II activity. The exact function of the repeats remains to be determined. Whether or not hClp1 is essential for the reconstitution of pre-mRNA cleavage could not be tested, as hPcf11 was insoluble in the absence of its partner protein.

The biological role of the RNA 5’ kinase activity of hClp1 remains unclear. We find that hClp1 maintains this activity when it forms a complex with hPcf11 and thus, presumably, in the 3’ processing complex. However, mice homozygous for the K127A point mutation in the kinase active site of Clp1 have no obvious defect in pre-mRNA processing; their severe neurological phenotype appears to be related to a tRNA processing defect (Hanada et al. 2013). The interpretation of phenotypes is complicated by the weak residual kinase activity of the K127A mutant (this work) and by its disruptive effect on the tRNA splicing endonuclease complex (Hanada et al. 2013; Karaca et al. 2014; Schaffer et al. 2014). Similarly, mutations in the yClp1 ATP binding site disturb the association with yPcf11 and even affect binding of yPcf11 to the Rna14-Rna15 complex (Holbein et al. 2011; Ghazy et al. 2012; Haddad et al. 2012). Nevertheless, we were able to isolate mutant hClp1 with no detectable kinase activity as a complex with hPcf11; the complex proved active in pre-mRNA cleavage. Thus, at least under the conditions of the *in vitro* reaction, hClp1 kinase activity is not required for RNA cleavage, in agreement with the situation in yeast (Ramirez et al. 2008). Not only is hClp1 found as a component of the tRNA splicing endonuclease (Paushkin et al. 2004; Weitzer and Martinez 2007), but knock-down of the endonuclease subunit SEN2 has been reported to inhibit 3’ processing of pre-mRNA (Paushkin et al. 2004). However, to our knowledge a tRNA endonuclease association has never been reported for hPcf11, and the endonuclease was not found in the CF II preparation of de Vries et al. (2000). Cleavage activity of CF II reconstituted in the absence of the tRNA endonuclease suggests that the enzyme is not directly involved in pre-mRNA 3’ processing. The caveat is that a crude fraction was used for the reconstitution assays.

Yeast Pcf11 and yClp1 are stably associated with Rna14 and Rna15, forming the CF IA complex (Amrani et al. 1997; Gross and Moore 2001; Gordon et al. 2011). Yeast Pcf11 is thought to interact mostly with Rna14 (Gordon et al. 2011), but also with a conserved C-terminal domain of Rna15 (Qu et al. 2007). As the human orthologs of Rna14 and Rna15 are CstF-77K and CstF-64K, an interaction between CF II and CstF might be expected. However, the interaction surface in yPcf11, amino acids 331-417 (Lionel Minvielle-Sebastia, personal communication), does not appear to be conserved in the mammalian orthologue, and we found no evidence for an interaction. On the basis of co-purification and pull-down from nuclear extract, an association of CF II with CF I was suggested (de Vries et al. 2000). Our experiments with highly purified recombinant preparations revealed no such association. In agreement with the data of Rüegsegger et al. (1996), the negative results of our interaction assays suggest that protein-protein interactions within the 3’ processing complex may be established mostly via CPSF. Indeed, isolated hClp1 bound CPSF in pull-down assays in nuclear extract (de Vries et al. 2000). Recombinant CPSF is not yet available in a form suitable for interaction assays with reconstituted CF II.

Our experiments revealed a high affinity of CF II for RNA. Binding is mediated predominantly by hPcf11. The length-dependence of CF II binding to a series of G homopolymers revealed that the protein requires ~ fourteen nucleotides for high affinity binding. A requirement for such an extended sequence is probably explained by the domain structure of hPcf11: The two zinc fingers provide the largest fraction of the total binding energy. Two zinc fingers can recognize nine nucleotides (Hudson et al. 2004). Binding of the remaining nucleotides must be mediated by more N-terminal regions of hPcf11, including the CID (Zhang et al. 2005; Hollingworth et al. 2006) and perhaps the highly charged region. An important role for RNA binding by Pcf11 is supported by the observation that mutations in the zinc fingers of yPcf11, in particular the second finger, strongly affect yeast growth and CF IA function in pre-mRNA processing (Guegueniat et al. 2017). CF II does not appear to have a very pronounced sequence-specificity of binding, but G-rich sequences were preferred. Not every polyadenylation site contains a G-rich sequence - for example the L3 RNA used in some of our experiments does not - but such motifs are statistically enriched far downstream of the cleavage site (see Introduction), and binding of yPcf11 has been mapped downstream of the cleavage site (Baejen et al. 2017). Thus, CF II might function in pre-mRNA processing via binding to these G-rich downstream elements. A role of CF II in the recognition of the pre-mRNA is consistent with the observation that knock-down of hPcf11 affects alternative polyadenylation (Li et al. 2015). Although the effects of hPcf11 knock-down were not strongly correlated with specific sequence motifs, several G-rich hexamers were overrepresented in the downstream region of proximal poly(A) sites that were downregulated by the knock- down, consistent with such hexamers being involved in hPcf11 binding. Members of the hnRNP H family of proteins have also been suggested to affect 3’ end processing via binding to downstream G-rich sequence elements (Bagga et al. 1998; Veraldi et al. 2001; Arhin et al. 2002). The relative contributions of these proteins and CF II to the recognition of the G-rich elements remain to be established.

## Materials and Methods

### Purification of His-Flag-tagged CF II

A stable HEK293 cell line expressing His-FLAG-tagged hClp1 (Paushkin et al. 2004) was obtained from Christopher Trotta. Affinity purification via FLAG beads followed by Ni^2+^ beads was carried out as described (Paushkin et al. 2004). After SDS-polyacrylamide gel electrophoresis, the main bands were identified by MS analysis (Paul Jenö, Biozentrum, University of Basel).

### Expression clones

Primers for the construction of *E. coli* expression plasmids are listed in Supplemental Table 2. All ORFs amplified by PCR were checked by sequencing.

Plasmids for *E. coli* expression of HisSumo-Pcf11zcz and HisSumo-Pcf11ΔN1184 were generated by amplification of the hPcf11 fragments and cloning into pETSumoAdapt (Bosse-Doenecke et al. 2008) with BsaI and NotI. For expression of HisSumo-CstF64 ΔN530 (aa 531-577), a PCR product was cloned into pET-SumoAdapt with BsaI and XhoI. A plasmid for GST-Pcf11zcz was made by cloning of a PCR fragment into pGEX6p1 (GE Healthcare) with BamHI and NotI.

ORFs coding for the following protein complexes were cloned into the Multibac system (Berger et al. 2004; Fitzgerald et al. 2006): CF II, composed of hClp1 and hPcf11, including mutant variants of both subunits; CstF, composed of CstF-77, CstF-64 and CstF- 50; CF I, composed of CF I-68 and CF I-25. All cDNAs are listed in Suppl. Table 3, transfer plasmids and resulting baculoviruses in Suppl. Table 4, and tags in Suppl. Table 5. Additional details, including primers used for cloning, will be provided upon request.

### Protein expression and purification

Protein purification was carried out under refrigeration. The pH of all buffers was adjusted at 5°C.

For CF II expression, Sf21 cells cultured in ExCell 420 serum-free medium (Sigma-Aldrich) at 27.5 °C and 125 rpm were infected with a baculovirus encoding hPcf11 and his-tagged hClp1 at an MOI ≥ 1. Cells were harvested ~72h after infection, washed with cold PBS, frozen in liquid N_2_ and stored at −80°C. Cells were thawed on ice, resuspended in lysis buffer (50 mM Tris-HCl, pH 8.5, 200 mM KCl, 0.2 mM EDTA, 0.02% NP-40, 10% sucrose, 1 mM DTT, 1 mM PMSF, 1 µg/ml each pepstatin and leupeptin) to ~1×10^7^ cells/ml and lysed by sonification. The lysate was clarified by centrifugation and applied to Ni-NTA agarose (Qiagen) (1 ml for 4 x 10^8^ cells starting material). The mixture was rotated for 2 h, packed into a column, washed with lysis buffer plus 10 mM imidazole, and protein was eluted with lysis buffer plus 500 mM imidazole. Fractions containing CF II were combined and slowly diluted with lysis buffer lacking KCl to a conductivity corresponding to 150 mM KCl. The solution (~ 5 mg protein) was applied to a 1 ml MonoQ column (GE Healthcare), which had been equilibrated in lysis buffer plus 150 mM KCl and was eluted with a gradient up to 1 M KCl in lysis buffer. Material from the MonoQ peak fractions (1.2 ml) was applied to a 100 ml Superose 6 column (GE Healthcare) equilibrated in 20 mM Tris-HCl, pH 8.5, 200 mM KCl, 0.1 mM EDTA, 10% sucrose, 1 mM DTT, 5 µM ZnCl_2_. Calibration proteins were ferritin (440 kDa, Stoke’s radius = 6.1 nm), catalase (232 kDa, 5.2 nm), BSA (66.4 kDa, 3.55 nm), ovalbumin (43 kDa, 3.0 nm) and RNase A (13.7 kDa, 1.64 nm). Fractions were analysed by SDS-polyacrylamide gel electrophoresis and Coomassie staining. CF II was quantified by comparison of hClp1 to a BSA standard curve. Peak fractions were combined and concentrated to ~1 µM. Gel filtration was omitted for most CFII variants containing hClp1 mutants. For CF II preparations destined to be used in cross-linking experiments, Tris was replaced by triethanolamine in all buffers. This had no effect on RNA binding or kinase activity.

His-tagged Clp1 was expressed and purified on Ni-NTA agarose as above. Material from the peak fraction (1.5 ml) was applied to a 120 ml Superdex 200 prep grade column (GE Healthcare) equilibrated in 20 mM Tris-HCl, pH 8.5, 200 mM KCl, 0.2 mM EDTA, 10% sucrose, 1 mM DTT, 0.1 mM ATP. The column was run and calibrated as above.

For expression of CstF, Sf21 cells were infected with a virus encoding all three subunits, including his-tagged CstF-77, and harvested as above. Cells were resuspended in 50 mM Tris-HCl, pH 8.0, 200 mM KCl, 0.5 mM EDTA, 0.02% NP-40, 10% glycerol, 1 mM DTT, 1 mM PMSF, 1 µg/ml pepstatin and leupeptin, and Ni-NTA purification was carried out as above. CstF-containing fractions (9 mg) were combined, dialysed against 50 mM Tris-HCl, pH 8.0, 75 mM KCl, 0.5 mM EDTA, 0.02% NP-40, 10 % glycerol, 0.5 mM DTT and applied to a 1 ml ResourceQ column (GE Healthcare) equilibrated with the same buffer. The column was eluted with a gradient to 1 M KCl in the same buffer. CstF-containing fractions were analyzed by SDS-PAGE, and protein content was quantified by Bradford assay.

CF I (including his-tagged CFI-25) was expressed in Sf21 cells as above. Cells were resuspended in 50 mM Tris-HCl, pH 8.0, 300 mM KCl, 0.5 mM EDTA, 10% sucrose, 1 mM DTT, 1 mM PMSF, 1 µg/ml pepstatin and leupeptin, and Ni-NTA purification was carried out as before. Eluted protein (~6 mg) was diluted to a conductivity equivalent to 20 mM Tris-HCl, pH 8.0, 170 mM KCl, 10% sucrose, 1 mM DTT and applied to a 1 ml ResourceQ column equilibrated in this buffer. The column was eluted and analyzed as above.

HisSumo-Pcf11ΔN1184, HisSumo-Pcf11zcz and GST-Pcf11zcz were expressed in *E. coli* Rosetta cells in TB medium plus 20 µM ZnCl_2_ with antibiotic selection. Expression was induced with 1 mM IPTG, and the culture was shaken overnight at 16-18°C. Cells were harvested and resuspended in lysis buffer (50 mM Tris-HCl, pH 8.0, 200 mM KCl, 10% sucrose). 20 µM ZnCl_2_ was included for Pcf11zcz purifications, and 10 mM imidazole was included for his-tagged proteins. A spatula tip of DNase I and lysozyme, 1 mM DTT and protease inhibitors (as above) were added, cells were broken in a French Press, and the lysate was cleared by centrifugation. All buffers used in the subsequent purification contained 1 mM DTT. HisSumo-Pcf11ΔN1184 from a 1500 ml culture was absorbed to 3 ml NiNTA agarose, which was then washed with 20 mM imidazole, 1 M KCl and again 20 mM imidazole, all in lysis buffer. Protein was eluted with lysis buffer plus 250 mM imidazole. Fractions containing the desired protein were pooled, mixed with Ulp1 Sumo protease (a kind gift of Bodo Moritz) and dialyzed against lysis buffer plus 20 mM imidazole overnight. The protein was then passed over a Ni-NTA column, and the flow-through and wash fractions were applied to a MonoQ column, which was eluted with a gradient to 1 M KCl in 50 mM Tris-HCl, pH 8.0, 10% sucrose. HisSumo-Pcf11zcz was purified in a similar manner except that fractions eluted from Ni-NTA were directly applied to the MonoQ column without cleavage of the HisSumo tag and the MonoQ buffer contained 20 µM ZnCl_2_. Lysate from a 1 l expression culture of GST-Pcf11zcz was applied to 1 ml of GSH-Sepharose (GE Healthcare), which was washed with 50 mM Tris-HCl, pH 8.0, 200 mM KCl, 10 % Sucrose, 2 mM DTT, 20 µM ZnCl2 and eluted with the same buffer plus 20 mM gluthathione. The protein was further purified by MonoQ chromatography as above.

Expression of HisSumo-CstFΔN530 in *E. coli* BL21 codon plus was induced by 1 mM IPTG in 800 ml SB-Medium containing kanamycin at 18°C overnight. Cells were broken in a French press and proteins bound to Ni-NTA beads in buffer A (50 mM Tris-HCl, pH 8.0, 300 mM KCl, 10% glycerol, 10 mM imidazole, 0.02% NP-40, 1 mM DTT) containing PMSF, leupeptin and lepstatin at 1 µg/ml each. The protein was eluted by buffer A plus 500 mM imidazole. Fractions containing His-Sumo-CstF64ΔN530 were pooled and dialysed against buffer A with 50 mM KCl and no imidazole.

Full-length bovine poly(A) polymerase has been described (Kühn et al. 2009).

### RNA

Cleavage substrate L3 and its ‘delta’ derivative with a point mutation in the AAUAAA sequence have been described (Humphrey et al. 1987). The SV40 late RNA containing the SV40 late 3’ processing signal was obtained by transcription of the −140/+70 DNA fragment (Conway and Wickens 1987) cloned into pSP64. The SV40 late Δ derivative had a U to G mutation in the polyadenylation signal. RNAs were made by in vitro transcription from DraI- linearized plasmids with SP6 RNA polymerase (Roche) under standard conditions in the presence of [α-^32^P]-UTP and cap analog (NEB). Uncapped SV40 late 40-mer fragments were made by *in vitro* transcription from DNA oligonucleotide templates (Supplemental Table 2) with T7 RNA polymerase (NEB) in the presence of [α-^32^P]-UTP and 20% DMSO. Synthetic RNAs were obtained from biomers.net GmbH (Ulm, Germany) and radiolabeled with polynucleotide kinase or with hClp1, which proved more efficient for some very G-rich oligonucleotides.

CD spectra were recorded in a J-815 Spectrometer (JASCO) between 340 nm and 220 nm at 1 cm path length. RNA concentration was 2 µM in 50 mM Tris-HCl, pH 8, 10% glycerol, 0.01% NP-40, 60 mM KCl, 40 mM NaCl at 20°C. Prior to measurement, the RNA was heated to 95°C for 5 min in 10 mM HEPES-NaOH, pH 7.0, and cooled on ice.

### Pre-mRNA cleavage assays

Cleavage assays contained 12.5 µl 2x cleavage buffer (40 mM HEPES-KOH, pH 8.0, 150 mM KCl, 4 mM DTT, 2 mM MgCl_2_, 40 mM creatine phosphate, 1 mM 3’-dATP, 7 % (w/v) polyethylene glycol 6000, 0.1 mg/ml tRNA), 60 fmol substrate RNA, and up to 8 µl protein fractions. The volume was made up to 25 µl with filter binding buffer (see below). Reactions were incubated at 30°C for up to 2 h and stopped by the addition of SDS-containing buffer. After proteinase K digestion, RNA was ethanol precipitated and analyzed by polyacrylamide -urea gel electrophoresis and phosphoimaging.

For a partial purification of cleavage factors, HeLa cell nuclear extract (Ipracell, Mons, Belgium) was mixed with ammonium sulfate (30 % saturation) and stirred on ice for at least one hour. After centrifugation, the pellet was dissolved in purification buffer (20 mM HEPES-KOH, pH 8.0, 3 mM MgCl_2_, 10 mM NaF, 0.2 mM EDTA, 0.5 mM DTT, 25% glycerol, 1x protease inhibitor cocktail [Biotools]) plus 300 mM KCl. An aliquot of 2.5 ml (13 mg protein) was fractionated over a 100 ml Superose 6 gel filtration column (GE Healthcare). This resulted in fractions that reconstituted pre-mRNA cleavage when 3 µl was supplemented with 100 fmol poly(A) polymerase and CF II. Active fractions were diluted to 80 mM KCl in purification buffer and passed over a Resource Q column (GE Healthcare), which was eluted with a gradient to 1 M KCl. Fractions were obtained that reconstituted cleavage when 1 µl was complemented with poly(A) polymerase, CF II and CstF at 50 - 100 fmol each. 50 - 100 fmol of CF I was also added, although the reaction did not depend on it.

### Kinase assays

Reactions were performed at 30°C in 50 mM HEPES-KOH, pH 8.0, 100 mM KCl, 5 mM MgAc, 5 µM ZnCl_2_, 10% sucrose, 0.02% NP-40, 1 mM DTT, 400 U/ml RNasin (Promega), C_14_ and ATP trace-labeled with ɣ-[^32^P]-ATP. For titrations of C_14_, ATP was used at 1 mM. The reaction mixture lacking ATP was pre-incubated for 5 min at 30°C and the reaction started by nucleotide addition. At different times, 5 µl aliquots were taken from the reaction, stopped with 5 µl 60 mM EDTA, 7 M urea and put on ice. Samples were heated for 3 min at 95°C and loaded on a denaturing 20% polyacrylamide gel. Phosphorylated C_14_ was quantified by phosphoimaging. For titrations of ATP, C_14_ was used at 20 µM. ATP contributed by the hClp1 column buffer was negligible. The reaction product was detected by adsorption to DEAE paper (Stayton and Kornberg 1983). Initial velocities were fitted to the Michaelis-Menten equation (SigmaPlot version 12.5). Kinase assays for the analysis of column fractions were done at 20 µM C_14_ and 0.625 or 1 mM ATP for 10 min.

### RNA binding assays

Nitrocellulose filter binding was done essentially as described (Kühn et al. 2003). RNA was heated at 95°C for 3 min in 10 mM HEPES-NaOH, pH 7.0, and cooled on ice before addition to the binding reaction. For the determination of apparent equilibrium dissociation constants, a fixed amount of RNA, typically 0.1 nM, was titrated with increasing amounts of protein. Data were fitted to a 1:1 association equilibrium with a single rectangular hyperbolic function (SigmaPlot version 12.5).

For electrophoretic mobility shift assays, binding reactions were set up and preincubated as for NC filter binding and analyzed on 5% (60:1) polyacrylamide gels run in 0.5x TBE at 8°C. Gels were dried on Whatman 3MM paper, and RNA was detected and quantified with phosphoimaging. Free and bound RNA were quantified as fractions of the total radioactivity in each lane; no-protein reactions were subtracted as background from the bound fraction. Apparent equilibrium dissociation constants were calculated as above.

UV cross-linking was carried out with 2 µM of 5’-labeled U_14_ RNA and 0.5 µM CF II in 50 µl 25 mM Tris-HCl, pH 8.5, 100 mM KCl, 10% sucrose, 0.02% NP-40, 0.2 mM EDTA, 0.5 mM ATP, 5 mM MgCl, 0.5 mM DTT. The mix was preincubated for 5 min at RT and UV-irradiated at 250 nm for 1 min (Kühn et al. 2003). Cross-linked products were analyzed without RNase digestion via SDS–polyacrylamide gel electrophoresis and phosphoimaging or used for immunoprecipitation: 5 mg of protein A Sepharose CL-4B (GE Healthcare) per sample was washed in 25 mM Tris-HCl, pH 8.5, 10% sucrose, 0.02% NP-40, 0.2 mM EDTA, 0.5 mM DTT, 200 mM KCl, mixed with 10 µl polyclonal α-Pcf11 antibody serum (made by Isabelle Kaufmann) or pre-immune serum and incubated for 1 h with end-over-end rotation at 8°C. Beads were washed three times with binding buffer and eluted with 4× SDS sample buffer for 5 min at 95°C. Cross-linked proteins were analyzed as above.

### RNA selection

100 nM hPcf11ΔN1184 in 400 µl filter binding buffer (25 mM Tris-HCl, pH 8.0, 50 mM KCl, 2 mM MgCl_2_, 0.01% NP-40, 1 mM DTT, 10% (v/v) glycerol, 0.1 U/µl Ribolock [ThermoFischer]) was mixed with 15 µg phosphorylated 14-mer RNA of the sequence N_14_GUUU and incubated for 30 min at room temperature with rotation. The mixture was filtered over a nitrocellulose membrane (GE Healthcare) equilibrated with filter binding buffer containing 15 µg/ml heparin, which was then washed with 20 ml ice-cold filter binding buffer. The spot where the binding reaction had been applied was cut out, and the RNA was eluted by addition of 250 µl 50% formamide, 1.8 M sodium acetate, 2 mM EDTA, 0.2% (w/v) SDS and shaking at 70°C for 30 min.

For full length CFII, 100 nM protein in 400 µl binding buffer (25 mM Tris-HCl, pH 7.5, 150 mM KCl, 3 mM MgCl_2_, 0.01% (v/v) NP-40, 1 mg/ml BSA, 5% (v/v) glycerol, 0.1 U/µl Ribolock) was mixed with 20µl IMAC sepharose beads (GE Healthcare) preloaded with Ni^2+^ ions and 15 µg of the N14GUUU RNA pool and incubated for 30 min at room temperature with agitation. Beads were collected by centrifugation (1000 g, 2 min, 4°C) and washed three times with ice-cold binding buffer. Bound RNA was eluted in 200 µl 10 mM Tris-HCl, pH 7.0, 400 mM NaCl, 1 mM EDTA, 1% (w/v) SDS.

For both types of experiments, no-protein controls were also performed. Selected RNA molecules were purified by phenol-chloroform extraction and ethanol precipitation and ligated with a 3´-DNA adaptor (AAACTGGAATTCTCGGGTGCCAAGG-Amino-C7) and a 5´- RNA adaptor containing a T7 promoter sequence (GUUCAGUAAUACGACUCACUAUAGGG). The products were reverse-transcribed with the First Strand cDNA Synthesis Kit (Thermo) and the primer

GCCTTGGCACCCGAGAATTCCAGTTT. PCR was used to amplify the cDNA sequence and introduce barcodes for next generation sequencing (NGS). PCR products were run on a 6% polyacrylamide urea gel, and the band at 150bp corresponding to the desired product was excised. The DNA was eluted overnight in 0.4 M NaCl and precipitated with ethanol, dissolved and stored for NGS analysis.

For the generation of RNA for a second round of selection, 50 ng of the PCR-pool was amplified with the primers AATGATACGGCGACCACCGAGATCTACACGTTCAGTAATACGACTCACTATAGG and GCCTTGGCACCCGAGAATTCCAGTTT. The PCR-product was purified with a PCR Clean-up Kit (Macherey-Nagel) and cleaved by addition of 1.5 µl FastDigest MssI (ThermoFisher), which recognizes the restriction site GTTTAAAC generated by ligation with the 3´adaptor. The cleaved DNA was transcribed with T7 polymerase, which yields a new pool of RNAs with the sequence GGG(N)_14_GUUU. The RNA was purified on an 18% polyacrylamide urea gel, dephosphorylated with FastAP (ThermoFisher) and phosphorylated with polynucleotide kinase. 15 µg of the prepared RNA were used in a second selection cycle, ligated and amplified as described above.

Libraries were sequenced on a MiSeq instrument (Illumina) with a 150 cycle MiSeq Reagent Kit to which we added a custom Read1 sequencing primer (5´- GATCTACACGTTCAGTAATACGACTCACTATAGGG-3´). Analysis was restricted to sequences from the second round of selection. Reads were barcode sorted and filtered for sequences containing the 3´ adaptor and the full Mss1-cleavage site, indicative of ligation of an intact RNA from the selection pool. After clipping of the adaptor and invariant sequence, only reads of the correct length (17 nt) were used for further analysis. The first three nucleotides were trimmed, and the resulting 14-mer sequences were analyzed for enriched sequence motifs using Weeder2 (Zambelli et al. 2014). Between 14,000 and 44,000 useful reads were obtained per experiment.

### Limited digestion of CF II

Per time point, 0.5 µg CF II was digested with 0.5 ng of sequencing-grade trypsin (Promega) in 15 µl 50 mM triethanolamine-HCl, pH 8.0, 200 mM KCl, 10% sucrose, 1 mM DTT at 16 °C. The reaction was stopped with 15 µl SDS loading buffer. For sequence analysis, 12.5 µg CF II was digested under comparable conditions. The reaction was stopped with 2 mM AEBSF, and the complex of his-tagged hClp1 and the trypsin-resistant hPcf11 fragment isolated on Ni-NTA agarose. The hPcf11 fragment was excised from an SDS-polyacrylamide gel, and fragments generated by trypsin or Asp-N digestion were analyzed by MS (Shevchenko et al. 2006). For Edman degradation, NiNTA-purified complex was separated by SDS-polyacrylamide gel electrophoresis and blotted to a PVDF membrane. After Ponceau S staining, the trypsin-resistant hPcf11 fragment was cut out and sequenced by the Proteome Factory AG (Berlin).

### Chemical cross-linking of CF II

For cross-linking, CF II was purified in buffers in which Tris was substituted by triethanolamine. 180 µl CF II (1 µM in column buffer) was mixed with 20 µl DSBU (12.8 mM in DMSO) (Müller et al. 2010) and incubated for 30 min at room temperature. The reaction was quenched with 200 µl of 2x SDS loading buffer containing Tris, and proteins were separated on a 4-12 % SDS-polyacrylamide gel. In-gel digestion of cross-linked protein complexes with trypsin, data acquisition and analysis were done as described (Götze et al. 2015; Arlt et al. 2016). Three gel bands of high molecular weight cross-linked complexes were analyzed separately by LC/MS/MS using three different collision energies (30% or 35% NCE or 29 +/- 5% stepped NCE). Cross-links identified at least twice at an FDR ≤ 1% were taken into account and manually validated. Data are available via ProteomeXchange with identifier PXD010177.

## Acknowledgements

We are grateful to Mathias Lorbeer for generating the CF I and CstF viruses and developing the first purification protocols; to Michael Götze for advice on data analysis of protein-protein cross-linking; to Christopher Trotta for the stable cell line expressing His-Flag-hClp1; to Paul Jenö for the analysis of purified His-Flag-hClp1; to Tobias Gruber for help with CD spectroscopy; to Bodo Moritz for the Ulp1 sumo protease; to Gudrun Scholz, Sandra Grund and Moritz Schmidt for help with cloning and protein purification; and to Lionel Minvielle-Sebastia for communicating unpublished results. This work was supported by grants from the Deutsche Forschungsgemeinschaft to EW (WA 548 15/1 and GRK 1591) and to GM (PP 1935).

## References

Amrani N, Minet M, Wyers F, Dufour M-E, Aggerbeck LP, Lacroute F. 1997. PCF11 encodes a third protein component of the yeast cleavage and polyadenylation factor I. Mol Cell Biol 17: 1102–1109.

Arhin GK, Boots M, Bagga PS, Milcarek C, Wilusz J. 2002. Downstream sequence elements with different affinities for the hnRNP H/H’ protein influence the processing efficiency of mammalian polyadenylation signals. Nucleic Acids Res 30: 1842–1850.

Arlt C, Gotze M, Ihling CH, Hage C, Schafer M, Sinz A. 2016. Integrated Workflow for Structural Proteomics Studies Based on Cross-Linking/Mass Spectrometry with an MS/MS Cleavable Cross-Linker. Anal Chem 88: 7930–7937.

Baejen C, Andreani J, Torkler P, Battaglia S, Schwalb B, Lidschreiber M, Maier KC, Boltendahl A, Rus P, Esslinger S et al. 2017. Genome-wide Analysis of RNA Polymerase II Termination at Protein-Coding Genes. Mol Cell 66: 38–49.

Bagga PS, Arhin GK, Wilusz J. 1998. DSEF-1 is a member of the hnRNP H family of RNA-binding proteins and stimulates pre-mRNA cleavage and polyadenylation in vitro. Nucleic Acids Res 26: 5343–5350.

Bagga PS, Ford LP, Chen F, Wilusz J. 1995. The G-Rich Auxiliary Downstream Element Has Distinct Sequence and Position Requirements and Mediates Efficient 3’ End Pre-Messenger-Rna Processing through a Trans-Acting Factor. Nucleic Acids Res 23: 1625–1631.

Barilla D, Lee BA, Proudfoot NJ. 2001. Cleavage/polyadenylation factor IA associates with the carboxyl-terminal domain of RNA polymerase II in Saccharomyces cerevisiae. Proc Natl Acad Sci U S A 98: 445–450.

Bedrat A, Lacroix L, Mergny JL. 2016. Re-evaluation of G-quadruplex propensity with G4Hunter. Nucleic Acids Res 44: 1746–1759.

Berger I, Fitzgerald DJ, Richmond TJ. 2004. Baculovirus expression system for heterologous multiprotein complexes. Nat Biotechnol 22: 1583–1587.

Beyer K, Dandekar T, Keller W. 1997. RNA ligands selected by cleavage stimulation factor contain distinct sequence motifs that function as downstream elements in 3’-end processing of pre-mRNA. J Biol Chem 272: 26769–26779.

Bienroth S, Wahle E, Suter-Crazzolara C, Keller W. 1991. Purification and characterisation of the cleavage and polyadenylation specificity factor involved in the 3’processing of messenger RNA precursors. J Biol Chem 266: 19768–19776.

Birse CE, Minvielle-Sebastia L, Lee BA, Keller W, Proudfoot NJ. 1998. Coupling termination of transcription to messenger RNA maturation in yeast. Science 280: 298–301.

Bosse-Doenecke E, Weininger U, Gopalswamy M, Balbach J, Knudsen SM, Rudolph R. 2008. High yield production of recombinant native and modified peptides exemplified by ligands for G-protein coupled receptors. Protein Expres Purif 58: 114–121.

Chan SL, Huppertz I, Yao C, Weng L, Moresco JJ, Yates JR, 3rd, Ule J, Manley JL, Shi Y. 2014. CPSF30 and Wdr33 directly bind to AAUAAA in mammalian mRNA 3’ processing. Genes Dev 28: 2370–2380.

Clerici M, Faini M, Aebersold R, Jinek M. 2017. Structural insights into the assembly and polyA signal recognition mechanism of the human CPSF complex. Elife 6: e33111.

Clerici M, Faini M, Muckenfuss LM, Aebersold R, Jinek M. 2018. Structural basis of AAUAAA polyadenylation signal recognition by the human CPSF complex. Nat Struct Mol Biol 25: 135–138.

Conway L, Wickens M. 1987. Analysis of mRNA 3’ end formation by modification interference: the only modifications which prevent processing lie in AAUAAA and the poly(A) site. EMBO J 6: 4177–4184.

Dalziel M, Nunes NM, Furger A. 2007. Two G-rich regulatory elements located adjacent to and 440 nucleotides downstream of the core poly(A) site of the intronless melanocortin receptor 1 gene are critical for efficient 3’ end processing. Mol Cell Biol 27: 1568–1580.

de Vries H, Rüegsegger U, Hübner W, Friedlein A, Langen H, Keller W. 2000. Human pre-mRNA cleavage factor IIm contains homologs of yeast proteins and bridges two other cleavage factors. EMBO J 19: 5895–5904.

Di Giammartino DC, Li W, Ogami K, Yashinskie JJ, Hoque M, Tian B, Manley JL. 2014. RBBP6 isoforms regulate the human polyadenylation machinery and modulate expression of mRNAs with AU-rich 3’ UTRs. Genes Dev 28: 2248–2260.

Dikfidan A, Loll B, Zeymer C, Magler I, Clausen T, Meinhart A. 2014. RNA specificity and regulation of catalysis in the eukaryotic polynucleotide kinase Clp1. Mol Cell 54: 975–986.

Eaton JD, Davidson L, Bauer DLV, Natsume T, Kanemaki MT, West S. 2018. Xrn2 accelerates termination by RNA polymerase II, which is underpinned by CPSF73 activity. Genes Dev 32: 127–139.

Erickson HP. 2009. Size and Shape of Protein Molecules at the Nanometer Level Determined by Sedimentation, Gel Filtration, and Electron Microscopy. Biol Proced Online 11: 32–51.

Fitzgerald DJ, Berger P, Schaffitzel C, Yamada K, Richmond TJ, Berger I. 2006. Protein complex expression by using multigene baculoviral vectors. Nat Methods 3: 1021–1032.

Ghazy MA, Gordon JM, Lee SD, Singh BN, Bohm A, Hampsey M, Moore C. 2012. The interaction of Pcf11 and Clp1 is needed for mRNA 3’-end formation and is modulated by amino acids in the ATP-binding site. Nucleic Acids Res 40: 1214–1225.

Gordon JM, Shikov S, Kuehner JN, Liriano M, Lee E, Stafford W, Poulsen MB, Harrison C, Moore C, Bohm A. 2011. Reconstitution of CF IA from overexpressed subunits reveals stoichiometry and provides insights into molecular topology. Biochem 50: 10203–10214.

Götze M, Pettelkau J, Fritzsche R, Ihling CH, Schäfer M, Sinz A. 2015. Automated Assignment of MS/MS Cleavable Cross-Links in Protein 3D-Structure Analysis. J Am Soc Mass Spectr 26: 83–97.

Gross S, Moore C. 2001. Five subunits are required for reconstitution of the cleavage and polyadenylation activities of Saccharomyces cerevisiae cleavage factor I. Proc Natl Acad Sci U S A 98: 6080–6085.

Grzechnik P, Gdula MR, Proudfoot NJ. 2015. Pcf11 orchestrates transcription termination pathways in yeast. Gene Dev 29: 849–861.

Guegueniat J, Dupin AF, Stojko J, Beaurepaire L, Cianferani S, Mackereth CD, Minvielle-Sebastia L, Fribourg S. 2017. Distinct roles of Pcf11 zinc-binding domains in pre- mRNA3’-end processing. Nucleic Acids Res 45: 10115–10131.

Guo AL, Gu HB, Zhou J, Mulhern D, Wang Y, Lee KA, Yang V, Aguiar M, Kornhauser J, Jia XY et al. 2014. Immunoaffinity Enrichment and Mass Spectrometry Analysis of Protein Methylation. Mol Cell Proteomics 13: 372–387.

Haddad R, Maurice F, Viphakone N, Voisinet-Hakil F, Fribourg S, Minvielle-Sebastia L. 2012. An essential role for Clp1 in assembly of polyadenylation complex CF IA and Pol II transcription termination. Nucleic Acids Res 40: 1226–1239.

Hallais M, Pontvianne F, Andersen PR, Clerici M, Lener D, Benbahouche NE, Gostan T, Vandermoere F, Robert MC, Cusack S et al. 2013. CBC-ARS2 stimulates 3’-end maturation of multiple RNA families and favors cap-proximal processing. Nature Struct Mol Biol 20: 1358–1366.

Hanada T, Weitzer S, Mair B, Bernreuther C, Wainger BJ, Ichida J, Hanada R, Orthofer M, Cronin SJ, Komnenovic V et al. 2013. CLP1 links tRNA metabolism to progressive motor-neuron loss. Nature 495: 474–480.

Holbein S, Scola S, Loll B, Dichtl BS, Hubner W, Meinhart A, Dichtl B. 2011. The P-Loop Domain of Yeast Clp1 Mediates Interactions Between CF IA and CPF Factors in Pre-mRNA 3’ End Formation. Plos One 6: e29139.

Hollingworth D, Noble CG, Taylor IA, Ramos A. 2006. RNA polymerase II CTD phosphopeptides compete with RNA for the interaction with Pcf11. RNA 12: 555–560.

Hu J, Lutz CS, Wilusz J, Tian B. 2005. Bioinformatic identification of candidate cis-regulatory elements involved in human mRNA polyadenylation. RNA 11: 1485–1493.

Hudson BP, Martinez-Yamout MA, Dyson HJ, Wright PE. 2004. Recognition of the mRNA AU-rich element by the zinc finger domain of TIS11d. Nature Struct Mol Biol 11: 257–264.

Humphrey T, Christofori G, Lucijanic V, Keller W. 1987. Cleavage and polyadenylation of messenger RNA precursors in vitro occurs with large and specific 3’ processing complexes. EMBO J 6: 4159–4168.

Hwang HW, Park CY, Goodarzi H, Fak JJ, Mele A, Moore MJ, Saito Y, Darnell RB. 2016. PAPERCLIP Identifies MicroRNA Targets and a Role of CstF64/64tau in Promoting Non-canonical poly(A) Site Usage. Cell Reports 15: 423–435.

Karaca E, Weitzer S, Pehlivan D, Shiraishi H, Gogakos T, Hanada T, Jhangiani SN, Wiszniewski W, Withers M, Campbell IM et al. 2014. Human CLP1 mutations alter tRNA biogenesis, affecting both peripheral and central nervous system function. Cell 157: 636–650.

Kaufmann I, Martin G, Friedlein A, Langen H, Keller W. 2004. Human Fip1 is a subunit of CPSF that binds to U-rich RNA elements and stimulates poly(A) polymerase. EMBO J 23: 616–626.

Kim DE, Chivian D, Baker D. 2004. Protein structure prediction and analysis using the Robetta server. Nucleic Acids Res 32: W526–W531.

Kim M, Vasiljeva L, Rando OJ, Zhelkovsky A, Moore C, Buratowski S. 2006. Distinct pathways for snoRNA and mRNA termination. Mol Cell 24: 723–734.

Kühn U, Buschmann J, Wahle E. 2017. The nuclear poly(A) binding protein of mammals, but not of fission yeast, participates in mRNA polyadenylation. RNA 23: 473–482.

Kühn U, Gündel M, Knoth A, Kerwitz Y, Rüdel S, Wahle E. 2009. Poly(A) tail length is controlled by the nuclear poly(A)-binding protein regulating the interaction between poly(A) polymerase and the cleavage and polyadenylation specificity factor. J Biol Chem 284: 22803–22814.

Kühn U, Nemeth A, Meyer S, Wahle E. 2003. The RNA binding domains of the nuclear poly(A) binding protein. J Biol Chem 278: 16916–16925.

Lackford B, Yao CG, Charles GM, Weng LJ, Zheng XF, Choi EA, Xie XH, Wan J, Xing Y, Freudenberg JM et al. 2014. Fip1 regulates mRNA alternative polyadenylation to promote stem cell self-renewal. EMBO J 33: 878–889.

Lambert NJ, Robertson AD, Burge CB. 2015. RNA Bind-n-Seq: Measuring the Binding Affinity Landscape of RNA-Binding Proteins. Method Enzymol 558: 465–493.

Li WC, You B, Hoque M, Zheng DH, Luo WT, Ji Z, Park JY, Gunderson SI, Kalsotra A, Manley JL et al. 2015. Systematic Profiling of Poly(A) plus Transcripts Modulated by Core 3’ End Processing and Splicing Factors Reveals Regulatory Rules of Alternative Cleavage and Polyadenylation. Plos Genet 11: e1005166.

Licatalosi DD, Geiger G, Minet M, Schroeder S, Cilli K, McNeil JB, Bentley DL. 2002. Functional interaction of yeast pre-mRNA 3’ end processing factors with RNA polymerase II. Mol Cell 9: 1101–1111.

Lunde BM, Reichow SL, Kim M, Suh H, Leeper TC, Yang F, Mutschler H, Buratowski S, Meinhart A, Varani G. 2010. Cooperative interaction of transcription termination factors with the RNA polymerase II C-terminal domain. Nature Struct Mol Biol 17: 1195-+.

Luo W, Johnson AW, Bentley DL. 2006. The role of Rat1 in coupling mRNA 3’-end processing to transcription termination: implications for a unified allosteric-torpedo model. Genes Dev 20: 954–965.

MacDonald CC, Wilusz J, Shenk T. 1994. The 64-kilodalton subunit of the CstF polyadenylation factor binds to pre-mRNAs downstream of the cleavage site and influences cleavage site location. Mol Cell Biol 14: 6647–6654.

Mandel CR, Kaneko S, Zhang HL, Gebauer D, Vethantham V, Manley JL, Tong L. 2006. Polyadenylation factor CPSF-73 is the pre-mRNA 3’-end-processing endonuclease. Nature 444: 953–956.

Martin G, Gruber AR, Keller W, Zavolan M. 2012. Genome-wide Analysis of Pre-mRNA 3’ End Processing Reveals a Decisive Role of Human Cleavage Factor I in the Regulation of 3’ UTR Length. Cell Reports 1: 753–763.

Mayer A, Schreieck A, Lidschreiber M, Leike K, Martin DE, Cramer P. 2012. The Spt5 C-Terminal Region Recruits Yeast 3’ RNA Cleavage Factor I. Mol Cell Biol 32: 1321–1331.

Meinhart A, Cramer P. 2004. Recognition of RNA polymerase II carboxy-terminal domain by 3’-RNA-processing factors. Nature 430: 223–226.

Minvielle-Sebastia L, Preker PJ, Wiederkehr T, Strahm Y, Keller W. 1997. The major yeast poly(A)-binding protein is associated with cleavate factor IA and functions in premessenger RNA 3’-end formation. Proc Natl Acad Sci USA 94: 7897–7902.

Müller MQ, Dreiocker F, Ihling CH, Schäfer M, Sinz A. 2010. Cleavable cross-linker for protein structure analysis: reliable identification of cross-linking products by tandem MS. Anal Chem 82: 6958–6968.

Noble CG, Beuth B, Taylor IA. 2007. Structure of a nucleotide-bound Clp1-Pcf11 polyadenylation factor. Nucleic Acids Res 35: 87–99.

Noble CG, Hollingworth D, Martin SR, Ennis-Adeniran V, Smerdon SJ, Kelly G, Taylor IA, Ramos A. 2005. Key features of the interaction between Pcf11 CID and RNA polymerase II CTD. Nat Struct Mol Biol 12: 144–151.

O’Reilly D, Kuznetsova OV, Laitem C, Zaborowska J, Dienstbier M, Murphy S. 2014. Human snRNA genes use polyadenylation factors to promote efficient transcription termination. Nucleic Acids Research 42: 264–275.

Oberg D, Fay J, Lambkin H, Schwartz S. 2005. A downstream polyadenylation element in human papillomavirus type 16 L2 encodes multiple GGG motifs and interacts with hnRNP H. J Virol 79: 9254–9269.

Paushkin SV, Patel M, Furia BS, Peltz SW, Trotta CR. 2004. Identification of a human endonuclease complex reveals a link between tRNA splicing and pre-mRNA 3’ end formation. Cell 117: 311–321.

Pearson E, Moore C. 2014. The evolutionarily conserved Pol II flap loop contributes to proper transcription termination on short yeast genes. Cell Rep 9: 821–828.

Porrua O, Libri D. 2015. Transcription termination and the control of the transcriptome: why, where and how to stop. Nat Rev Mol Cell Biol 16: 190–202.

Qian ZW, Wilusz J. 1991. An Rna-Binding Protein Specifically Interacts with a Functionally Important Domain of the Downstream Element of the Simian Virus-40 Late Polyadenylation Signal. Mol Cell Biol 11: 5312–5320.

Qu XP, Perez-Canadillas JM, Agrawal S, De Baecke J, Cheng HL, Varani G, Moore C. 2007. The C-terminal domains of vertebrate CstF-64 and its yeast orthologue Rna15 form a new structure critical for mRNA 3’-end processing. J Biol Chem 282: 2101–2115.

Ramirez A, Shuman S, Schwer B. 2008. Human RNA 5’-kinase (hClp1) can function as a tRNA splicing enzyme in vivo. RNA 14: 1737–1745.

Rüegsegger U, Beyer K, Keller W. 1996. Purification and characterization of human cleavage factor I-m, involved in the 3’ end processing of messenger RNA precursors. J Biol Chem 271: 6107–6113.

Rüegsegger U, Blank D, Keller W. 1998. Human pre-mRNA cleavage factor Im is related to spliceosomal SR proteins and can be reconstituted from recombinant subunits. Mol Cell 1: 243–253.

Sadofsky M, Connelly S, Manley JL, Alwine JC. 1985. Identification of a Sequence Element on the 3’ Side of Aauaaa Which Is Necessary for Simian-Virus 40 Late Messenger-Rna 3’-End Processing. Mol Cell Biol 5: 2713–2719.

Sadowski M, Dichtl B, Hubner W, Keller W. 2003. Independent functions of yeast Pcf11p in pre-mRNA 3’ end processing and in transcription termination. EMBO J 22: 2167–2177.

Schaffer AE, Eggens VR, Caglayan AO, Reuter MS, Scott E, Coufal NG, Silhavy JL, Xue Y, Kayserili H, Yasuno K et al. 2014. CLP1 founder mutation links tRNA splicing and maturation to cerebellar development and neurodegeneration. Cell 157: 651–663.

Schönemann L, Kühn U, Martin G, Schäfer P, Gruber AR, Keller W, Zavolan M, Wahle E. 2014. Reconstitution of CPSF active in polyadenylation: Recognition of the polyadenylation signal by WDR33. Genes Dev 28: 2381–2393.

Sheets MD, Stephenson P, Wickens MP. 1987. Products of in vitro cleavage and polyadenylation of simian virus 40 late pre-mRNAs. Mol Cell Biol 7: 1518–1529.

Shevchenko A, Tomas H, Havlis J, Olsen JV, Mann M. 2006. In-gel digestion for mass spectrometric characterization of proteins and proteomes. Nat Protoc 1: 2856–2860.

Shi Y, Di Giammartino DC, Taylor D, Sarkeshik A, Rice WJ, Yates JR, Frank J, Manley JL. 2009. Molecular architecture of the human pre-mRNA 3’ processing complex. Mol Cell 33: 365–376.

Shi Y, Manley JL. 2015. The end of the message: multiple protein-RNA interactions define the mRNA polyadenylation site. Genes Dev 29: 889–897.

Siegel LM, Monty KJ. 1966. Determination of Molecular Weights and Frictional Ratios of Proteins in Impure Systems by Use of Gel Filtration and Density Gradient Centrifugation. Application to Crude Preparations of Sulfite and Hydroxylamine Reductases. Biochim Biophys Acta 112: 346–362.

Stayton M, Kornberg A. 1983. Complexes of Escherichia coli primase with the replication origin of G4 phage DNA. J Biol Chem 258: 13205–13212.

Stojko J, Dupin A, Chaignepain S, Beaurepaire L, Vallet-Courbin A, Van Dorsselaer A, Schmitter JM, Minvielle-Sebastia L, Fribourg S, Cianferani S. 2017. Structural characterization of the yeast CF IA complex through a combination of mass spectrometry approaches. Int J Mass Spectrom 420: 57–66.

Sun Y, Zhang Y, Hamilton K, Manley JL, Shi Y, Walz T, Tong L. 2018. Molecular basis for the recognition of the human AAUAAA polyadenylation signal. Proc Natl Acad Sci U S A 115: E1419–E1428.

Takagaki Y, Manley JL. 1994. A polyadenylation factor subunit is the human homologue of the Drosophila suppressor of forked protein. Nature 372: 471–474.

Takagaki Y, Manley JL. 1997. RNA recognition by the human polyadenylation factor CstF. Mol Cell Biol 17: 3907–3914.

Venkataraman K, Brown KM, Gilmartin GM. 2005. Analysis of a noncanonical poly(A) site reveals a trinartite mechanism for vertebrate poly(A) site recognition. Genes Dev 19: 1315–1327.

Veraldi KL, Arhin GK, Martincic K, Chung-Ganster LH, Wilusz J, Milcarek C. 2001. hnRNP F influences binding of a 64-kilodalton subunit of cleavage stimulation factor to mRNA precursors in mouse B cells. Mol Cell Biol 21: 1228–1238.

Volanakis A, Kamieniarz-Gdula K, Schlackow M, Proudfoot NJ. 2017. WNK1 kinase and the termination factor PCF11 connect nuclear mRNA export with transcription. Genes Dev 31: 2175–2185.

Wang XY, Goodrich J, Gooding AR, Naeem H, Archer S, Paucek RD, Youmans DT, Cech TR, Davidovich C. 2017. Targeting of Polycomb Repressive Complex 2 to RNA by Short Repeats of Consecutive Guanines. Mol Cell 65: 1056–1067.

Waterhouse AM, Procter JB, Martin DMA, Clamp M, Barton GJ. 2009. Jalview Version 2-a multiple sequence alignment editor and analysis workbench. Bioinformatics 25: 1189–1191.

Weitzer S, Hanada T, Penninger JM, Martinez J. 2015. CLP1 as a novel player in linking tRNA splicing to neurodegenerative disorders. Wires RNA 6: 47–63.

Weitzer S, Martinez J. 2007. The human RNA kinase hClp1 is active on 3’ transfer RNA exons and short interfering RNAs. Nature 447: 222–226.

West S, Proudfoot NJ. 2008. Human Pcf11 enhances degradation of RNA polymerase II-associated nascent RNA and transcriptional termination. Nucleic Acids Research 36(3): 905–914.

Wittmann S, Renner M, Watts BR, Adams O, Huseyin M, Baejen C, El Omari K, Kilchert C, Heo DH, Kecman T et al. 2017. The conserved protein Seb1 drives transcription termination by binding RNA polymerase II and nascent RNA. Nat Commun 8.

Xiang K, Tong L, Manley JL. 2014. Delineating the structural blueprint of the pre-mRNA 3’- end processing machinery. Mol Cell Biol 34(11): 1894–1910.

Xu XQ, Perebaskine N, Minvielle-Sebastia L, Fribourg S, Mackereth CD. 2015. Chemical shift assignments of a new folded domain from yeast Pcf11. Biomol Nmr Assign 9(2): 421–425.

Yang F, Hsu P, Lee SD, Yang W, Hoskinson D, Xu W, Moore C, Varani G. 2017. The C terminus of Pcf11 forms a novel zinc-finger structure that plays an essential role in mRNA 3’-end processing. RNA 23: 98–107.

Yang Q, Coseno M, Gilmartin GM, Doublie S. 2011. Crystal Structure of a Human Cleavage Factor CFI(m)25/CFI(m)68/RNA Complex Provides an Insight into Poly(A) Site Recognition and RNA Looping. Structure 19(3): 368–377.

Yang W, Hsu PL, Yang F, Song J-E, Varani G. 2018. Reconstitution of the CstF complex unveils a regulatory role for CstF-50 in recognition of 3’-end processing signals. Nucleic Acids Res 46: 493–503.

Yonaha M, Proudfoot NJ. 1999. Specific transcriptional pausing activates polyadenylation in a coupled in vitro system. Molecular Cell 3(5): 593–600.

Zambelli F, Pesole G, Pavesi G. 2014. Using Weeder, Pscan, and PscanChIP for the Discovery of Enriched Transcription Factor Binding Site Motifs in Nucleotide Sequences. Curr Protocol Bioinformatics 47: 2.11.11–12.11.31.

Zhang ZQ, Fu JH, Gilmour DS. 2005. CTD-dependent dismantling of the RNA polymerase II elongation complex by the pre-mRNA 3’-end processing factor, Pcf11. Gene Dev 19(13): 1572–1580.

Zhang ZQ, Gilmour DS. 2006. Pcf11 is a termination factor in Drosophila that dismantles the elongation complex by bridging the CTD of RNA polymerase II to the nascent transcript. Molecular Cell 21(1): 65–74.

Zhu Y, Wang X, Forouzmand E, Jeong J, Qiao F, Sowd GA, Engelman AN, Xie X, Hertel KJ, Shi Y. 2018. Molecular mechanisms for CFIm-mediated regulation of mRNA alternative polyadenylation. Mol Cell 69: 62–74.

